# Designing smart spatial omics experiments with S2Omics

**DOI:** 10.1101/2025.09.21.677634

**Authors:** Musu Yuan, Kaitian Jin, Hanying Yan, Amelia Schroeder, Chunyu Luo, Sicong Yao, Bernhard Domoulin, Jonathan Levinsohn, Tianhao Luo, Jean R. Clemenceau, Inyeop Jang, Minji Kim, Yunhe Liu, Minghua Deng, Emma E. Furth, Parker Wilson, Anupma Nayak, Idania Lubo, Luisa Maren Solis Soto, Linghua Wang, Jeong Hwan Park, Katalin Susztak, Tae Hyun Hwang, Mingyao Li

**Author notes:** Correspondence: Mingyao Li.

## Abstract

Spatial omics technologies have transformed biomedical research by enabling high-resolution molecular profiling while preserving the native tissue architecture. These advances provide unprecedented insights into tissue structure and function. However, the high cost and time-intensive nature of spatial omics experiments necessitate careful experimental design, particularly in selecting regions of interest (ROIs) from large tissue sections. Currently, ROI selection is performed manually, which introduces subjectivity, inconsistency, and a lack of reproducibility. Previous studies have shown strong correlations between spatial molecular patterns and histological features, suggesting that readily available and cost-effective histology images can be leveraged to guide spatial omics experiments. Here, we present S2Omics, an end-to-end workflow that automatically selects ROIs from histology images with the goal of maximizing molecular information content in the ROIs. Through comprehensive evaluations across multiple spatial omics platforms and tissue types, we demonstrate that S2Omics enables systematic and reproducible ROI selection and enhances the robustness and impact of downstream biological discovery.

## Introduction

Recent advances in spatial omics technologies, particularly in spatial transcriptomics and proteomics^1, 2^, have enabled high-dimensional molecular profiling with preserved spatial context, offering unprecedented insights into tissue organization and disease pathogenesis. Despite these significant advancements, current commercial platforms remain expensive and often have limited tissue capture areas. For instance, Visium HD^3^ which provides subcellular resolution and whole-transcriptome coverage, costs about $7,000 per sample for a tissue capture area of only 6.5 mm × 6.5 mm, considerably smaller than typical tissue specimens. Other spatial omics platforms, including both spatial transcriptomics^4–9^ and spatial proteomics^10–14^, are similarly priced and constrained by small tissue capture areas. These technical and economic constraints necessitate careful experimental design. To avoid wasting valuable resources on suboptimal data acquisition, researchers must strategically select regions of interest (ROIs) within large tissue sections to ensure that spatial profiling captures the most biologically informative regions while minimizing cost and time investment.

Currently, ROI selection is predominantly manual, subjective, and non-reproducible, relying heavily on the expertise of pathologists. This process typically involves visual inspection of hematoxylin and eosin (H&E) stained histology images, which are widely accessible and cost-effective. However, because ROI selection is based on tissue morphology, the process is labor-intensive and inherently subjective, making it prone to human error. Such variability introduces inconsistencies across experiments and laboratories, poses significant challenges to reproducibility, and compromises the reliability of spatial omics studies.

The critical problem of ROI selection in spatial omics studies remains underexplored in the existing literature. While several recent efforts have aimed to improve experimental design in spatial omics, they do not address the core challenge of ROI selection within tissue sections. For instance, Jones *et al*.^15^ developed an algorithm to optimize tissue sectioning strategies, but their method does not consider ROI selection within a given section. SOFisher^16^ employs reinforcement learning for automated field-of-view (FOV) selection. Its sequential decision-making approach requires gene expression data from FOVs selected in the previous steps. This iterative process is impractical in real-world settings due to prolonged processing time and risk of RNA degradation. As such, these methods do not provide a solution to the practical challenge of ROI selection.

Here, we present S2Omics, a framework that addresses this critical methodological gap by providing a systematic, reproducible, and computationally efficient approach for ROI selection using only H&E histology images. Previous studies^17–20^ have shown that similar histological patterns often correspond to similar spatial molecular profiles, supporting the premise that H&E-based ROI selection can effectively capture molecular heterogeneity across tissues. We demonstrate the versatility of S2Omics across multiple tissue types, including human breast, colon, kidney, liver, and stomach. By applying S2Omics to experimental design in three leading spatial omics platforms, including Xenium, Visium HD, and CosMx, we validate that S2Omics consistently identified ROIs enriched with biologically informative patterns, often matching or outperforming manual selections made by pathologists. Moreover, we highlight that suboptimal ROI selection can compromise biological discovery, resulting in missed molecular signals. S2Omics fulfills essential criteria for real-world deployment, including objectivity, consistency, cost-effectiveness, and computational scalability. By standardizing the ROI selection step, S2Omics ensures that the substantial investment in spatial omics experiments yield maximally informative data, thereby establishing it as a foundational tool for the field.

## Results

### Overview of S2Omics

**Fig. 1** illustrates the S2Omics workflow, which consists of three components: histology image feature extraction, ROI selection, and whole slide molecular information recovery. To preserve tissue integrity for spatial molecular analysis on the target slice, S2Omics performs ROI selection using H&E image obtained from the immediately adjacent tissue section. The high architectural similarity between sequential thin tissue slices ensures that ROIs identified from the immediately adjacent H&E-stained section can effectively guide experimental design for the target slice. To characterize tissue architecture, S2Omics leverages pre-trained pathology image foundation models to extract both global and local image features for all superpixels of size 8 μm × 8 μm on the adjacent H&E image. S2Omics employs the UNI^21^ model by default, but is compatible with other foundation models^22–27^.

**Fig. 1.**
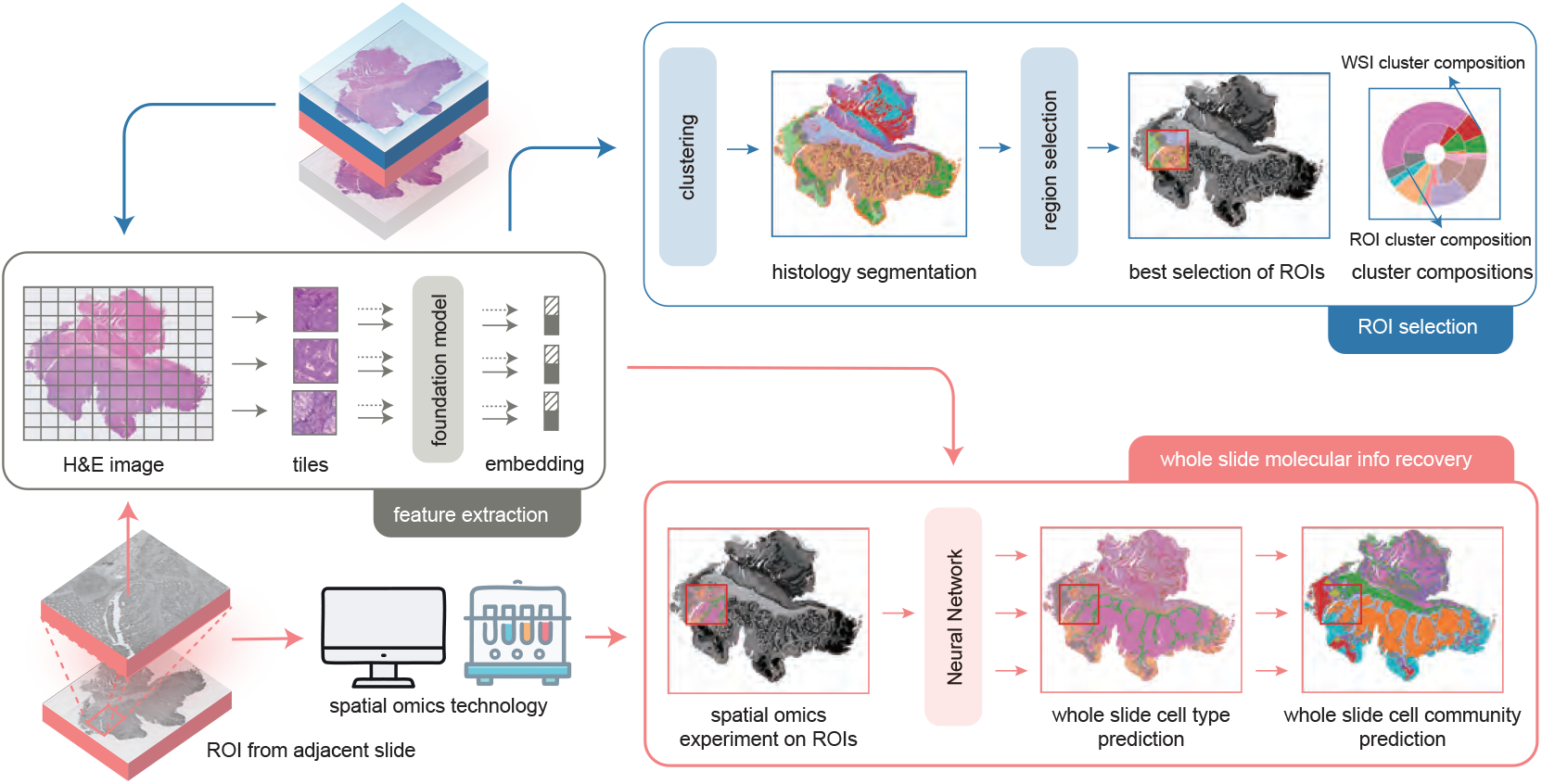
Workflow, ROI selection, and cell label broadcasting of S2Omics. Model summary of S2Omics. To preserve tissue integrity for spatial molecular analysis, S2Omics performs ROI selection using H&E image obtained from an adjacent tissue slice of the intended spatial omics section. The adjacent H&E image is divided into tiles and converted into hierarchical histology image features using foundation model. These features are used to cluster histology image into functionally diverse regions, i.e. histology clusters. ROI selection is then conducted with the clustering results to maximize ROI score. After the spatial omics experiments over the selected ROI is completed, cell type and cell community predicting models are trained to predict cell type and cell community labels in unsampled tissue regions.

The second component, ROI selection, is designed to identify ROIs that best capture the diversity of tissue structures present within a tissue section. The image features extracted in the first component are used to segment the tissue into histologically distinct regions via unsupervised clustering. S2Omics then samples candidate regions across the whole slide. We introduce a novel metric, the ROI score (see Methods), to quantitively assess the representativeness of each candidate region. Guided by this score, S2Omics selects ROIs that aim to maximize the molecular information content and minimize the experimental cost of spatial omics profiling.

The final component, whole slide molecular information recovery, is an optional virtual prediction module designed primarily to guide subsequent experiments, but it can also be used to validate ROI selection made by S2Omics. By propagating cell type and cell community labels from experimentally profiled ROIs across the entire tissue section using histological features, this module provides a tissue-wide perspective that informs downstream analyses and helps prioritize future experiments. In benchmarking contexts, the accuracy of these virtual predictions further serves as a biologically meaningful check that the selected ROIs are sufficiently representative to capture key molecular variation.

### Application to a human gastric cancer sample to guide Xenium experiment

To evaluate the performance of S2Omics, we applied it to a gastric cancer sample (~10 mm × 24 mm) that included both high-resolution H&E image and 10x Xenium spatial transcriptomics data. This sample was chosen for its relatively large size, the presence of diverse and interesting tissue structures, and the availability of single-cell resolution gene expression data spanning the entire tissue section. These characteristics allowed us to evaluate whether the ROIs identified by S2Omics could effectively capture gene expression variation across the whole tissue. As shown in **Fig. 2a**, the tissue slice included not only typical gastric structures, such as mucosa and intestinal metaplasia, but also tumor cells, and multiple tertiary lymphoid structures (TLSs). TLSs are of particular interest because they are positively associated with improved responses to immunotherapy and better clinical outcomes^28^.

**Fig. 2.**
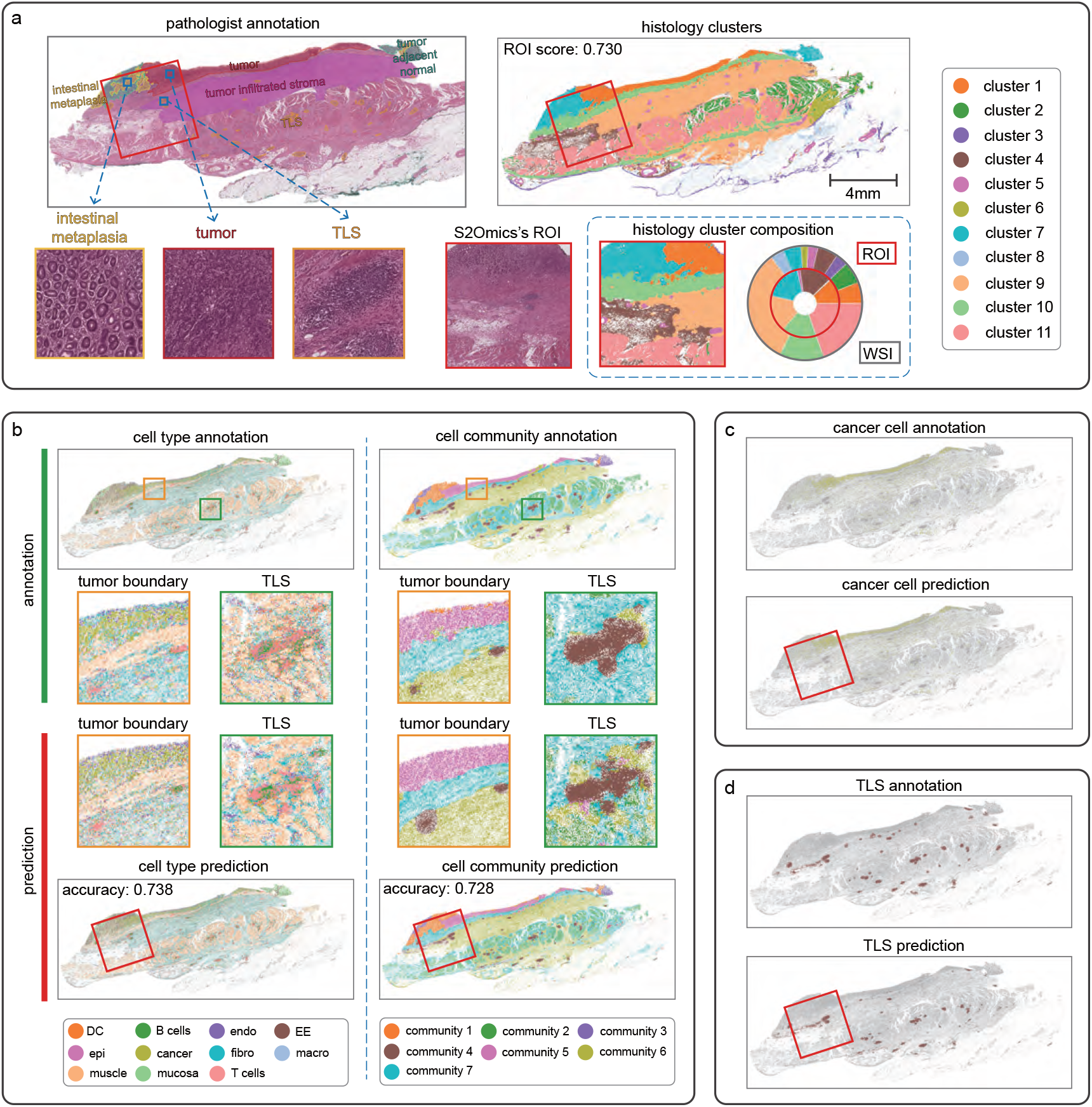
Application to a gastric cancer sample to guide Xenium experiment. **a**, ROI selected by S2Omics, visualized on H&E image and histology clusters; comparison between histology cluster compositions of ROI and whole-slide image (WSI). **b, c, d**, Evaluation of cell type and cell community label prediction. The Xenium data-based annotation served as ground truth for both cell type and cell community label prediction. Only the labels of superpixels inside the selected ROI were used for broadcasting model training to mimic the situation that spatial omics experiment was only conducted in the ROI. **b**, Visual comparison between the ground truth and S2Omics’s cell type and cell community predictions from the selected ROI. The ground truth and predicted image of an unsampled region at tumor boundary and another unsampled region involving a mature TLS are zoomed in for detailed comparison. **c**, Visual comparison between ground truth and S2Omics predicted cancer cells. **d**, Visual comparison between ground truth and S2Omics predicted TLSs.

We considered a scenario where the budget permits the measurement of only a single ROI. With a fixed ROI size of 4 mm × 4 mm, S2Omics identified an optimal ROI, highlighted by the red box in **Fig. 2a**. Detailed examination revealed that this ROI encompasses all key structures, including intestinal metaplasia, tumor, tumor adjacent normal, tumor infiltrated stroma, and TLS. Our histology segmentation results demonstrated strong concordance with pathologist annotation, showing significant overlaps between cluster 1 and tumor region, cluster 7 and intestinal metaplasia/tumor adjacent normal, cluster 9 and tumor-infiltrated stroma, among others. To further evaluate the selected ROI, we compared the histological cluster composition within the ROI (red, inner ring) to that of the entire tissue slice (grey, outer ring). Of the 11 clusters present in the tissue, seven were represented within the ROI, maintaining a balanced distribution. The four unsampled clusters corresponded to muscle region (clusters 2 and 6) and low-quality tissue regions (clusters 3 and 8). Although the clusters were identified solely based on histological information, they effectively captured meaningful cellular characteristics (**Supplementary Fig. 1a**). These findings indicate that, despite its smaller size, the selected ROI preserves the overall tissue architecture while enriched for underrepresented clusters.

To further examine if the ROI selected by S2Omics contains representative cell types and tissue structures, we broadcasted cell type and cell community labels from the ROI to the entire tissue slice and compared these predictions with annotations derived directly from Xenium gene expression measurements (**Fig. 2b**). Cell communities, which characterize distinct tissue structures, were identified using the methodology described in the Methods section. The cell type composition of each cell community is shown in **Supplementary Fig. 1b**. Overall, the broadcasting performance was satisfactory, achieving prediction accuracies of 73.8% for cell types and 72.8% for cell communities. These high accuracy values confirm that S2Omics selected ROIs contain sufficient cellular diversity to support robust inference of cellular and structural features from histological patterns. This validates both the biological relevance and representativeness of the automatically selected ROIs.

To further assess S2Omics’s performance, we focused on a tumor region and a TLS for detailed examination (**Fig. 2b**). In the tumor region, both the cell type and cell community predictions successfully recovered the tumor (community 5) and the tumor-infiltrated stroma (community 6, community 7) boundary with high precision. For the selected TLS, the predicted cell community labels were highly accurate, and the cell type predictions revealed the expected TLS structure of B cell aggregates surrounded by T cells. To comprehensively evaluate S2Omics’s ability to predict tumor cells and TLSs, we isolated both annotated and S2Omics predicted cancer cells and TLSs, and displayed the prediction results alongside the ground truth annotations (**Fig. 2c,d**). Notably, 73.4% of the tumor cells were correctly predicted. Among the 40 TLSs in the tissue, four were directly captured within the ROI, and 28 of the remaining 36 were recovered by S2Omics. Detailed true positive and false positive predictions for tumor cells and TLSs are shown in **Supplementary Fig.2a**, and a Sankey diagram illustrating the relationship between predicted cell level labels and ground truth annotations is provided in **Supplementary Fig. 2b**. Collectively, these results demonstrate that the ROI selected by S2Omics effectively identified representative tissue structures. Moreover, the broadcasting based on the spatial gene expression data measured in the ROI reliably expanded cell-level labels across the entire tissue slice, providing valuable information for subsequent experiments.

To quantitatively assess the performance of S2Omics, we conducted a systematic benchmarking analysis using 500 randomly selected 4 mm × 4 mm ROIs centered within the tissue slice. The ROI selected by S2Omics achieved the highest ROI score (0.73), ranking within the top three for cell type prediction accuracy and within the top ten for cell community prediction accuracy among all randomly selected ROIs (**Supplementary Fig. 2c**). It is reassuring that, despite relying solely on histology without molecular input, S2Omics identified a near-optimal ROI for spatial omics profiling, as reflected in its strong predictive performance. To further demonstrate the importance of ROI selection, we randomly selected an ROI with a mediocre ROI score (0.65) (**Supplementary Fig. 3a**). As demonstrated in **Supplementary Fig. 3b**, this ROI captured very few enteroendocrine (EE) cells and cells from communities 1, 2, 3, and 5. Consequently, the molecular and morphological characteristics of these populations could not be adequately characterized, resulting in decreased accuracy for cell-level label prediction (**Supplementary Fig. 3c**). This example illustrates how suboptimal ROI selection can lead to missed biological signals and compromise discovery potential, underscoring the critical importance of strategic ROI selection.

### Application to human colon cancer samples to guide Visium HD experiment

We next evaluated S2Omics by comparing its performance with expert’s ROI selection using a colorectal cancer section (P1 CRC, 20 mm × 16 mm). This section was measured using Visium HD within a 6.5 mm × 6.5 mm ROI selected by 10x Genomics researchers. This expert-selected ROI encompassed four distinct tissue compartments: invasive cancer, immune-infiltrated stroma, intestinal gland and smooth muscle. To ensure a fair comparison, we applied S2Omics under the constraint that only one ROI of size 6.5 mm × 6.5 mm could be selected within the given budget. The ROI selected by S2Omics (**Fig. 3a**, solid red box) showed remarkable concordance with the ROI chosen by the 10x Genomics researchers (**Fig. 3a**, dashed red box), covering 89.3% of expert-selected cells (superpixels within the pathologist’s ROI that passed quality control) while including 16.3% more valid superpixels. We carefully compared the histology cluster compositions of the two ROIs by visualizing them as pie charts (**Fig. 3b**). Compared to the expert’s selection, the ROI selected by S2Omics minimized blank space and achieved a more balanced distribution of histology clusters. Notably, it captured a higher proportion of the invasive cancer region, which is of particular interest to oncology researchers.

**Fig. 3.**
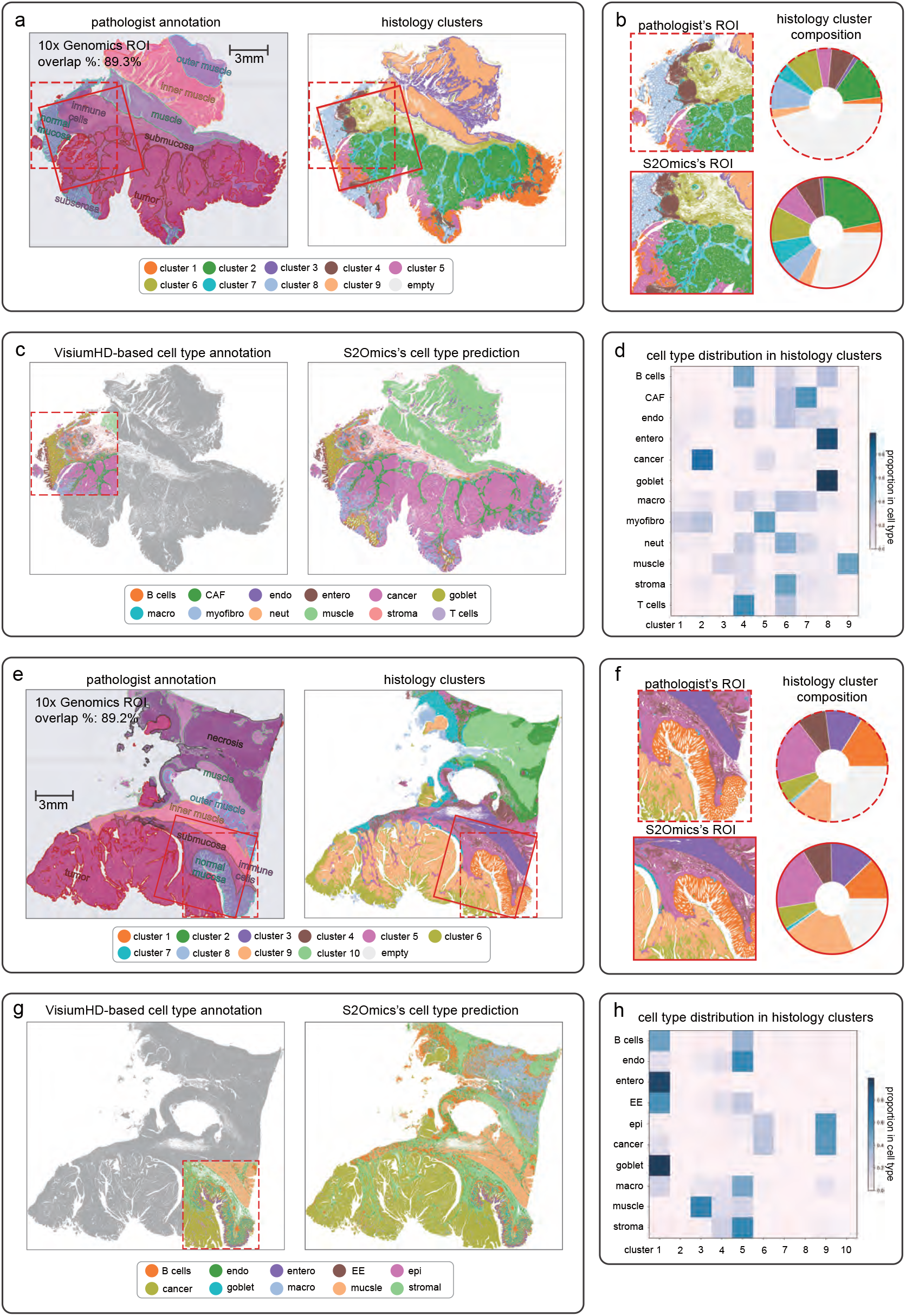
Application to two colorectal cancer samples to guide Visium HD experiment. **a**, Visual comparison between ROIs separately selected by S2Omics (red box) and experienced pathologist (dashed red box) on a colorectal cancer tissue section (P1 CRC). **b**, Histology cluster compositions of the two ROIs and their pie chart visualizations. **c**, Visium HD-based cell type annotation and S2Omics’s cell type prediction on the whole tissue section. For all superpixels that have Visium HD-based cell type annotation, their histology features and annotations were served as training data. Cell type labels of all superpixels that passed quality control were predicted using the trained model. **d**, Distribution of cell types in histology clusters. Each square shows the percentage of cells in corresponding cell type category and histology cluster occupied in all cells in that cell type category. **e, f, g, h**, Application and evaluation on another colorectal cancer sample with paired Visium HD data (P5 CRC).

Ideally, we would further evaluate the quality of S2Omics selected ROI by performing cell level label broadcasting with spatial gene expression data from the ROI. However, since such data are unavailable, we utilized the Visium HD data obtained from the ROI selected by 10x Genomics researchers. Given the 89.3% coverage of expert-selected cells, we expect the resulting performance to reflect the representativeness of samples captured by S2Omics’s ROI. As shown in **Fig. 3c,d**, the predicted cell type distributions from S2Omics closely aligned with pathologist’s annotation, demonstrating strong correspondence between cell types and tissue compartments. For example, malignant cells and CAFs corresponded to ‘tumor’ (cluster 1, cluster 2, cluster 5, cluster 7), B cells and T cells matched ‘immune cells’ (clusters 4 and 6), goblet cells aligned with ‘normal mucosa’ (cluster 8), and smooth muscle cells matched ‘muscle’ (clusters 3 and 9). This strong concordance demonstrates that S2Omics-selected ROIs capture sufficient representation of each cell type, enabling comprehensive biological characterization and accurate reconstruction of the spatial cellular distribution across the entire tissue specimen.

To further evaluate S2Omics’s performance, we applied it to two additional colon cancer samples and one healthy colon sample from different patients (**Fig. 3e-h, Supplementary Figs. 4,5**). As expected, the S2Omics selected ROIs largely overlap with the ROIs selected by 10x Genomics researchers, and the whole-slice cell type predictions generated by S2Omics were consistent with the pathologist’s annotations. Specifically, S2Omics’s ROI selection covered 89.2% of the superpixels in pathologist’s ROI for P5 CRC (**Fig. 3e-h**), 76.7% for P2 CRC (**Supplementary Fig. 4**), and 95.7% for P3 NAT (**Supplementary Fig. 5**). Notably, in P2 CRC, S2Omics’s predictions uncovered previously uncharacterized tissue structures, including a significant aggregation of B cells and T cells at the tumor boundary in the lower-left region of the section (**Supplementary Fig. 4e**). This structure, which was not captured within the original Visium HD ROI, highlights S2Omics’s utility beyond ROI selection. By providing researchers with a detailed preview of tissue architecture across unmeasured tissue regions, S2Omics offers valuable insights that can guide decisions on additional spatial omics experiments and help identify biologically important regions for further investigation.

### Application to human kidney samples to guide CosMx experiment

Encouraged by the promising performance of S2Omics in guiding Xenium and Visium HD experiments, we next applied it to another popular spatial transcriptomics platform, CosMx^29^, which involves FOV selection. Unlike other spatial transcriptomics technologies, CosMx experiments are conducted on small FOVs (0.5 mm × 0.5 mm each) rather than larger ROIs. The experiment’s runtime increases with the number of FOVs, creating a tradeoff between the area captured and data quality due to RNA degradation over time. In most cases, the number of FOVs in a single CosMx slide (~15 mm × 20 mm) is limited to 200, covering approximately 1/6 of the chip area. Exceeding this limit often results in the average number of molecules per cell dropping below 100, which is inadequate for downstream analysis. However, even with a limited number of FOVs, many cells remain unidentified due to the low capture rate. These practical constrains underscore the critical importance of optimizing FOV selection to balance tissue coverage and data quality.

The data analyzed included two kidney tissue sections: one from a healthy sample (left in **Fig. 4a**) and the other from a sample with type 2 diabetes (T2D) (right in **Fig. 4a**). H&E image-based tissue segmentation revealed sample heterogeneity attributable to differences in disease status, but also identified common tissue structure, such as glomeruli (cluster 3). Given the unique characteristics of CosMx experiments, we developed two modes for experimental design. The first mode identifies optimal FOVs, which may be non-contiguous, while the second mode selects larger ROIs (5 mm × 5 mm), suitable for researchers interested in obtaining a contiguous set of FOVs within each ROI. Both the FOV and ROI selections made by S2Omics for these two samples are shown in **Fig. 4a**, where the numbers of FOVs and ROIs were automatically determined as described in the Methods section. When applying S2Omics, we considered the two tissue samples simultaneously as they were positioned on the same slide, and the H&E image encompassed both. Notably, the FOVs and ROIs selected by S2Omics effectively covered both samples. A closer examination of the histology composition pie charts in **Fig. 4a** revealed that S2Omics balanced the selection across different functional regions in both the FOV and ROI modes, given equal importance to both tissue sections.

**Fig. 4.**
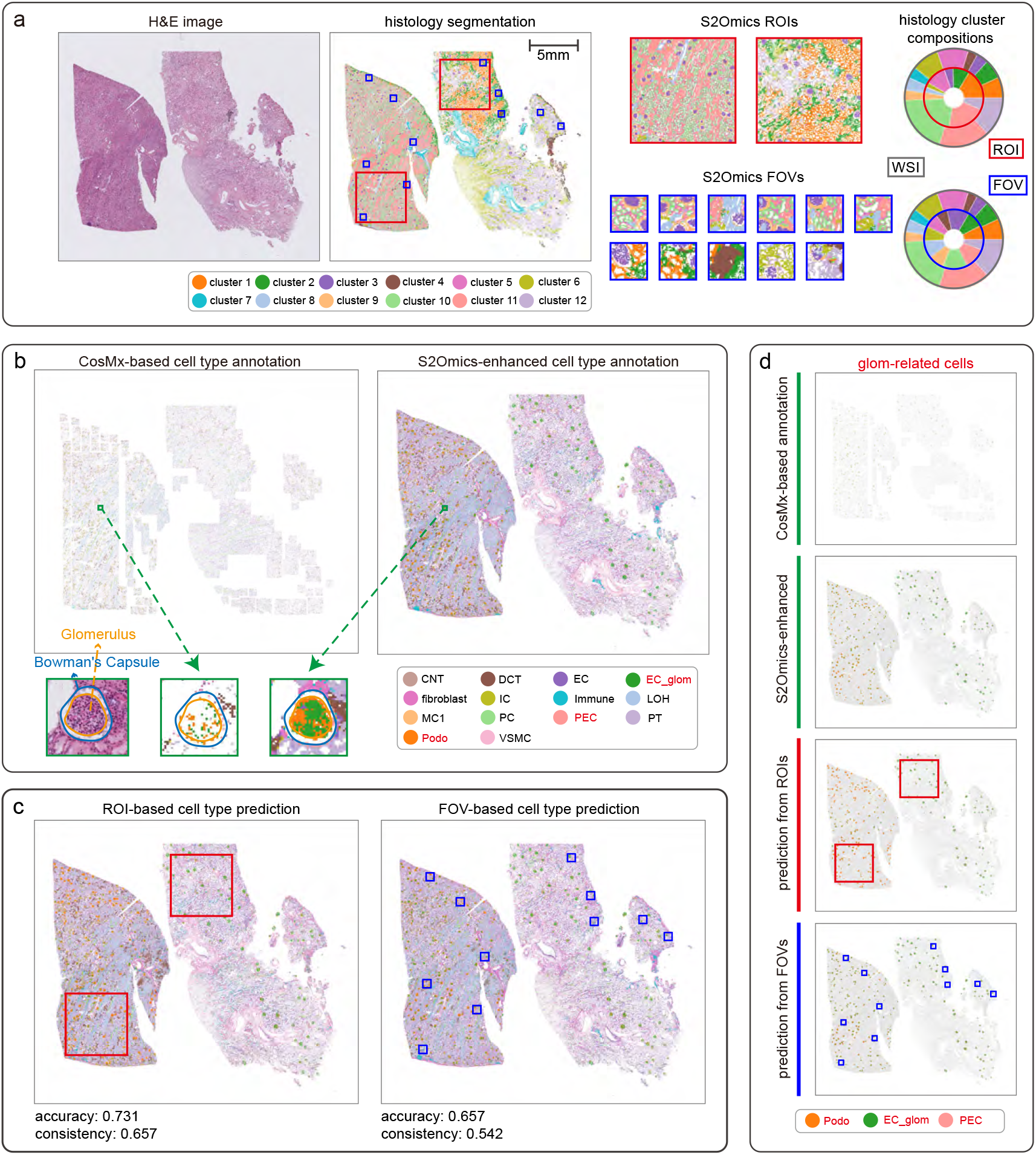
Application to healthy and diabetes kidney samples to guide and enhance CosMx experiment. **a**, ROIs and FOVs selected by S2Omics; comparison between histology cluster compositions of ROIs and WSI; comparison between histology cluster compositions of FOVs and WSI. The size of each FOV is 0.5 mm × 0.5 mm, the real physical size of CosMx FOVs. And ROI size is 5 mm × 5 mm. **b**, Visual comparison between CosMx-based cell type annotation and S2Omics’s cell type prediction. For all superpixels that have CosMx-based cell type annotation, their histology features and annotations were served as training data. Cell type labels of all superpixels that passed quality control were predicted using the trained model through their histology features. **c**, Visualization of S2Omics’s cell type predictions based on ROIs and FOVs. The prediction models were trained only using spatial omics data in the ROIs or the FOVs. **d**, Visual comparison between glomerulus-related cells identified with CosMx data and glomerulus-related cells predicted by S2Omics. The glomerulus-related cells include glomerular endothelial cells (EC_glom), podocytes (podo) and glomerular parietal epithelial cells (PEC.)

To further access the quality of S2Omics selected ROIs, we performed cell level label broadcasting. However, due to the limited tissue coverage in the original CosMx data and the low cell and molecule capture efficiency of the measured FOVs, the data exhibited patchy patterns with significant tissue gaps. As a result, gene expression was missing for many cells, making it infeasible to assign gene expression-based cell level labels as gold standard for evaluation. To address this issue, we first annotated cell type labels for cells captured in the CosMx experiment and then used S2Omics to broadcast these labels to the surrounding, unmeasured cells. The resulting CosMx-based and S2Omics-enhanced cell type distributions are shown in **Fig. 4b**. The dense and contiguous predictions generated by S2Omics enabled clearer identification of tissue structures, as demonstrated by a glomerulus surrounded by Bowman’s capsule.

With both CosMx based and the enhanced cell type labels, we assessed the quality of S2Omics’s ROIs/FOVs through cell-type broadcasting experiments using two key metrics: prediction accuracy compared to the CosMx derived labels and prediction consistency compared to the enhanced labels. As shown in **Fig. 4c**, when the broadcasting model was trained using CosMx data from the S2Omics selected ROIs, we achieved a cell type prediction accuracy of 0.731 and a consistency of 0.657. Even when trained on just 11 small FOVs, the model maintained relatively high accuracy (0.657) and consistency (0.542). Although the accuracy for FOV based recovery was lower, the two ROIs covered a significantly larger tissue area compared to the 11 small FOVs (50 mm^2^ vs. 2.75 mm^2^). This stark difference demonstrates that S2Omics can efficiently capture representative cells from a wide range of cell types and tissue structures and enable robust feature characterization even under strict budget constraints. Since RNA and protein degradation occur over time often forces spatial omics experiments to balance quality against tissue coverage, S2Omics offers a practical solution. Its intelligent FOV selection strategy maximizes the capture of biologically informative regions while minimizing the total experimental area required, thereby helping to mitigate the quality-versus-scale trade-off.

Optimal ROI selection should preserve not only cellular diversity but also key tissue structures. In the kidney, one such critical structure is the glomerulus, which comprises three key cell types: glomerular endothelial cells (EC_glom), podocytes (podo), and glomerular parietal epithelial cells (PEC). Previous studies have established that T2D is associated with a reduction in podocyte abundance, contributing to glomerular loss in diabetic nephropathy^30, 31^. Therefore, an effective ROI selection strategy in this context should capture both healthy and T2D glomeruli while enabling robust identification of associated cell types using cells within the selected regions. As shown in **Fig. 4a**, S2Omics selected one ROI from the healthy tissue and one from the T2D tissue. For FOV-based selection, six FOVs were identified in the healthy section and five in the T2D section. Both selection strategies successfully captured representative glomeruli from each condition, allowing accurate reconstruction of the spatial distribution of glomerular cell types across the entire tissue specimen. **Fig. 4d** shows that, using only cell type labels derived from the selected ROIs or FOVs, S2Omics precisely reconstructed the spatial architecture of glomeruli, effectively capturing differences between healthy and T2D tissues. These results highlight S2Omics’s ability to identify and preserve distinct, biologically relevant structures, even when such features exhibit subtle morphological differences. Moreover, S2Omics implicitly addresses common technical challenges in spatial transcriptomics, such as low cell capture rates and missing FOVs, by generating dense, contiguous, and whole-slide cell type maps. This significantly enhanced both the quality and interpretability of CosMx data, facilitating more robust downstream biological investigations.

### Incorporating prior knowledge in ROI selection

In the previous applications, S2Omics treated all tissue regions equally during ROI selection, which is suitable for most scenarios. However, we recognize the importance of allowing user-defined prioritization of specific tissue regions of interest. To demonstrate S2Omics’s ability to incorporate prior knowledge in ROI selection, we analyzed a colorectal cancer tissue section (P1 CRC), where smooth muscle tissue that occupies about 32.1% of the tissue section was of lower interest (**Fig. 5a**, red contour). Without prior information, S2Omics’s default selection (**Fig. 5b**) included 5.1% of muscle associated regions (cluster 3, cluster 9). When negative prior information about smooth muscle was incorporated into ROI selection (**Fig. 5a**, red superpixels), S2Omics identified a better ROI than the default selection, ignoring all the smooth muscle region.

**Fig. 5.**
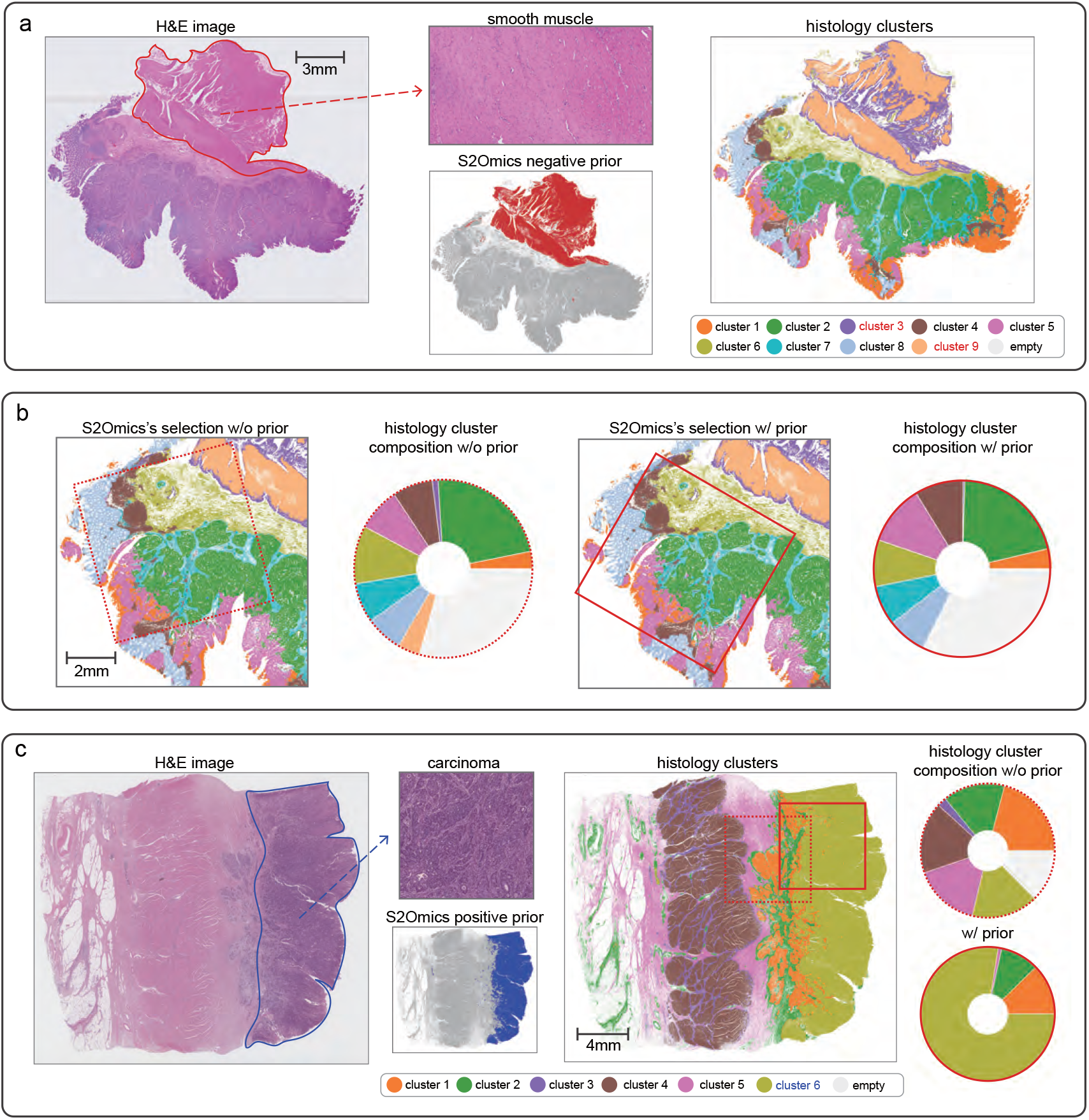
Incorporation of prior knowledge to tailor ROI selection. **a, b**, an example of incorporating ‘negative prior’ information in ROI selection, applying S2Omics on a colorectal cancer tissue section (P1 CRC). ‘Negative prior’ refers to regions that are of less interest to researchers and can be discarded in ROI selection. In contrast, ‘positive prior’ refers to regions that researchers are mostly interested in. **a**, Visualization of ‘negative prior’ where ‘negative prior’ is smooth muscle dominated region. **b**, Visualization of histology cluster compositions for ROI selected by S2Omics without prior (dotted red box) and ROI selected by S2Omics with prior (red box). **c**, An example of incorporating ‘positive prior’ in ROI selection, applying S2Omics on a gastric cancer tissue section. Here, ‘positive prior’ is carcinoma region. Visualization of histology cluster compositions of S2Omics’s selected ROI without prior (dotted red box) and with prior (red box).

In practical applications, researchers may often have a specific interest in certain tissue areas, e.g., tumor region in a cancer sample. To illustrate this, we provided an example of adding ‘positive prior’ for a gastric cancer tissue section where tumor cell occupied approximately one third of the section (**Fig. 5c**, blue contour). The default S2Omics’s ROI selection algorithm identified a ROI that included 83.9% normal and 16.1% tumor regions, reflecting the overall normal and tumor cell distribution in the entire tissue section. To mimic a researcher’s preference for tumor focused analysis, we introduced a ‘positive prior’ by giving cluster 6, which matched well with carcinoma area, greater weight in the ROI selection process. This guided approach produced an ROI with 78.1% tumor content, demonstrating how user-defined priorities can effectively influence ROI selection and maximize the retrieval of desired information. In this example, the prior preference parameter α was set to 5, indicating the emphasized histology clusters are 5 times more important than other clusters (see **Methods** for details). Users can adjust this parameter to control the degree to which ROI selection prioritizes histology clusters with positive priors.

### Applications to additional tissue types

To demonstrate S2Omics’s broad applicability, we applied it to additional tissue types. First, we analyzed a breast cancer section (BC S1) with paired Xenium gene expression data to illustrate S2Omics’s ability to automatically determine the optimal number of ROIs. As shown in **Fig. 6a**, this sample includes both invasive cancer (cluster 9) and ductal carcinoma in situ (DCIS, cluster 8) regions. The invasive cancer is mainly located in the left half of the section, while the DCIS occupies the right half, making it challenging for a single small ROI to capture representative areas for both cancer types. To show a single ROI is insufficient to encompass all relevant tissue structures, we fixed the size of each ROI at 2 mm × 2 mm. **Fig 6a** displays the ROIs selected by S2Omics when the number of ROIs was set to one, two, and three. S2Omics determined that two 2 mm × 2 mm ROIs were optimal for this sample, as the ROI score increased significantly when increasing from one to two ROIs but only showed a minor improvement when expanding from two to three.

**Fig. 6.**
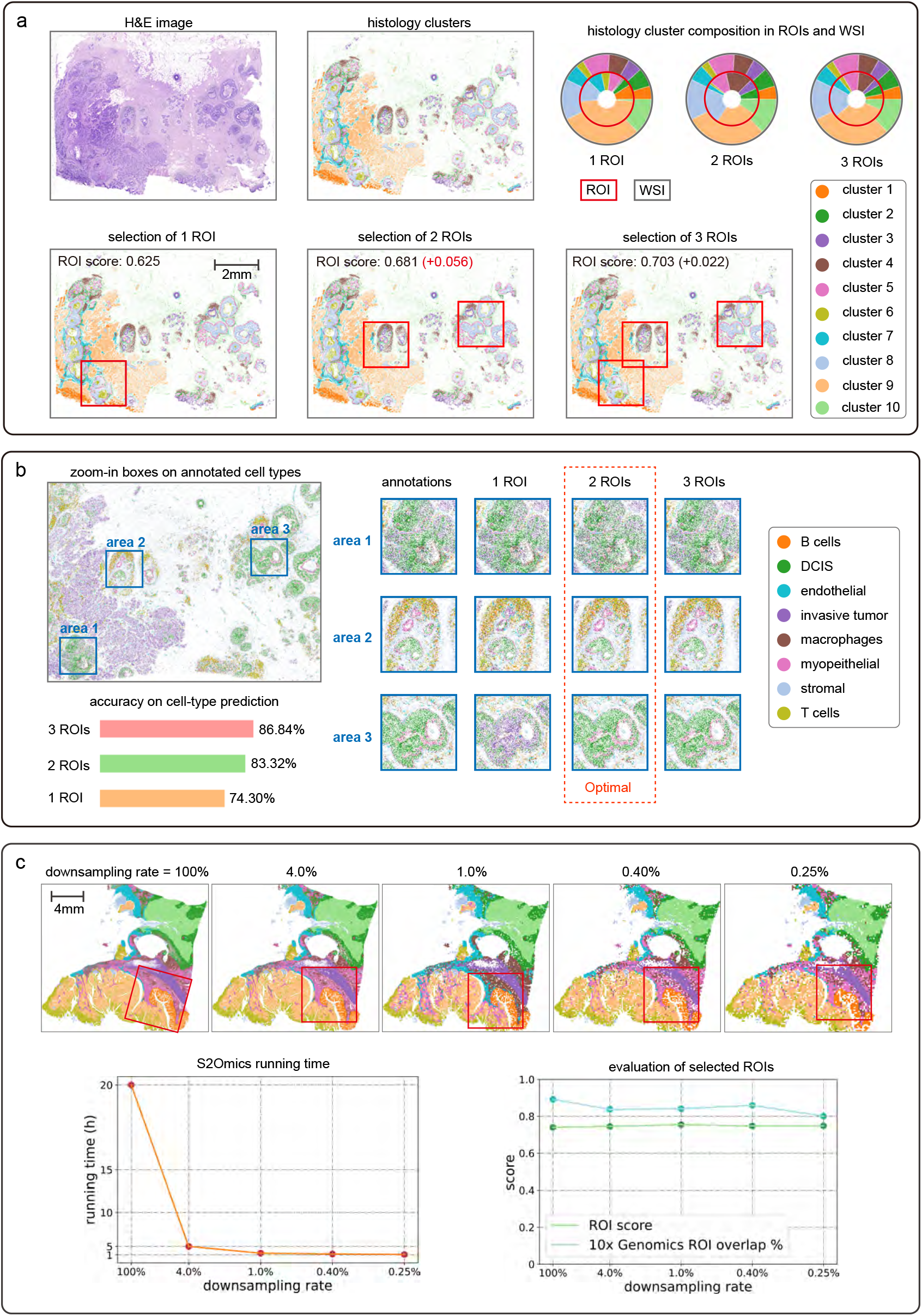
Automatic determination of optimal number of ROIs and running time evaluation. **a, b**, Application of S2Omics to a breast cancer tissue section with fixed ROI size (2mm × 2mm) but different numbers of ROIs. **a**, Visual comparison among S2Omics selected ROIs when the numbers of ROIs are 1, 2, 3, and 4, and corresponding pie for about histology cluster compositions. **b**, Xenium-based cell type annotation and cell type predictions using various numbers of ROIs. Three representative areas of this section were selected for detailed comparison, where ‘area 1’ is a DCIS region surrounded by invasive cancer region, ‘area 2’ is an immune-infiltrated DCIS, region and ‘area 3’ is a typical DCIS region. **c**, Comparison of running time and ROI quality when applying S2Omics to a colorectal cancer section (P5 CRC) with different down-sampling rates for histological feature extraction. The size of ROI was fixed at 6.5 mm × 6.5 mm. ROI quality was measured by ROI score and overlap rate with ROI selected by experienced pathologist.

To validate that two ROIs are sufficient while one ROI is not enough to effectively capture molecular variations in the tissue, we further examined cell type predictions from models trained using one, two, and three ROIs. For a more detailed comparison, we selected three small representative areas and analyzed the differences between Xenium derived cell type annotations and predictions obtained with different numbers of ROIs for model training. The selected areas include a DCIS region surrounded by invasive cancer (area 1), an immune-infiltrated DCIS (area 2), and a typical DCIS (area 3). As shown in **Fig. 6b**, all three models accurately predicted area 1. However, models trained with two and three ROIs provided better predictions for cell type distribution in areas 2 and 3 compared to the model trained with only one ROI. Notably, there was little difference between predictions obtained with two and three ROIs. The prediction accuracies are 75.1% with one ROI, 85.5% with two ROIs, 88.9% with three ROIs, and 90.6% with four ROIs. These results indicate that two ROIs are sufficient to effectively characterize the cellular variations and tissue microenvironments present in this tissue section, making it an optimal selection for this section. These findings further validate S2Omics’s ability to automatically determine the optimal number of ROIs for efficient and informative tissue profiling.

To demonstrate S2Omics’s flexibility in handling diverse tissue types, we conducted additional experiments in which the number of ROIs was automatically determined. We applied S2Omics to additional cancer tissue samples (**Supplementary Figs. 6-8**), and in all cases, it consistently selected optimal ROIs that captured representative tumor regions and enabled accurate reconstruction of spatial architecture across the entire tissue section. In a second breast cancer sample (BC S2) with paired Xenium gene expression data (**Supplementary Fig. 6**), S2Omics selected ROI led to 76.2% accuracy in cell type prediction, 53.8% accuracy in cell community prediction, and 90.6% accuracy in tumor cell prediction. Similar results were obtained in a liver cancer sample (**Supplementary Fig. 7**) and a kidney cancer sample (**Supplementary Fig. 8**). We further evaluated S2Omics on one healthy kidney section and one liver tissue section, both with paired Xenium gene expression data, to assess its ability to select ROIs that capture repetitive structural patterns. As shown in **Supplementary Figs. 9**,**10**, S2Omics effectively identified glomerular structure in the kidney and hepatic lobule structure in liver using minimal ROI areas. The biological information extracted from these ROIs enabled accurate reconstruction of cell types and cell communities across unmeasured tissue regions, highlighting the S2Omic’s potential to reduce experimental costs while preserving comprehensive biological information.

### Evaluation of S2Omics in non-standard scenarios

Previous evaluations focused on standard settings where tissues were large or histology is strongly correlated with molecular variation. To assess performance in more complex, real-world contexts, we examined four non-standard and more challenging scenarios. The first involved a whole-slide image containing multiple small breast cancer biopsies of varying quality, including distorted adipose-dominated regions. Pathologist annotations identified four tissue regions: invasive cancer, benign breast tissue, benign stroma, and benign fibro-adipose tissue (**Supplementary Fig. 11** left). Using 1 mm × 1 mm ROI specifications, S2Omics automatically selected two ROIs, with one from invasive cancer and one from mixed benign breast tissue and stroma, while avoiding selecting low-quality benign fibro-adipose (**Supplementary Fig. 11** right). These results reflected unbiased ROI selection without prior knowledge. With invasive cancer and histological clusters 1, 10, 13, and 14 specified as positive priors, S2Omics reoptimized ROI selection and identified two alternative ROIs that specifically targeted the invasive cancer regions, demonstrating adaptability to research-specific priorities (**Supplementary Fig. 12**). Beyond square ROIs, S2Omics also supports circular ROI selection, making it suitable for design of tissue microarrays (**Supplementary Fig. 13**).

Next, we applied S2Omics to three consecutive H&E-stained breast cancer sections^32^ containing both invasive cancer and immune infiltrates. Joint image segmentation of all sections yielded 15 consistent histological clusters, from which S2Omics selected a single ROI spanning with histological pattern spanning all sections and capturing both invasive cancer and immune infiltrates (**Supplementary Fig. 14**). This example demonstrates S2Omics’s capacity for robust, biologically relevant ROI selection in multi-section analyses.

In the third scenario, we analyzed a colorectal cancer dataset involving two consecutive H&E sections, one high-quality and one post-Xenium with staining artifacts (**Supplementary Fig. 15a**).

Despite severe batch effects between the two H&E images, S2Omics extracted histological features from both sections and performed joint segmentation, yielding 15 consistent clusters (**Supplementary Fig. 15b**). It then selected a representative ROI with histological patterns spanning both sections, spatially proximal to the ROI manually chosen by the 10x expert (**Supplementary Fig. 15c**). Additionally, cell type labels from Visium HD profiling of the 10x selected ROI in section 1 was successfully broadcast across both sections, demonstrating that label broadcasting effectively mitigates batch effects and supports multi-section analyses.

Finally, we tested S2Omics when histology and molecular profiles are weakly correlated. In a gastric cancer section with rare signet-ring cells (1.94% of spots) and subtle morphology, S2Omics segmented the H&E image into 15 clusters, with clusters 8 and 10 corresponding to the pathologist annotated signet-ring cells (**Supplementary Fig. 16**) and consistently enriching them across different ROI sizes (**Supplementary Fig. 17**). Importantly, signet-ring cells represent an early-stage diffuse-type gastric carcinoma feature that is difficult to detect from H&E alone, highlighting the challenge of this case. In breast cancer, where CD4 ^+^ and CD8^+^T cells are morphologically indistinguishable in H&E, S2Omics preserved their near-original proportions in selected ROIs (**Supplementary Fig. 18**). These results underscore S2Omics’s ability to identify informative ROIs even when morphological cues are weak.

### Robustness of S2Omics to the number of histology clusters and ROI size

S2Omics adopts a two-stage clustering strategy in which a tissue section is first partitioned into a relatively large number of preliminary clusters, followed by a guided merging process to produce the final segmentation (see Methods). Rather than requiring users to define an arbitrary similarity threshold for cluster merging, S2Omics allows users to directly specify the desired number of final clusters. To support this process, we perform hierarchical clustering analysis on the preliminary clusters and provide visualization of inter-cluster similarities. This enables users to identify cluster pairs with significantly higher similarity and merge them accordingly, ensuring biologically meaningful consolidation. To evaluate the impact of cluster number selection on ROI identification, we applied S2Omics to a gastric cancer dataset while varying the final cluster numbers from 5 to 20. Remarkably, S2Omics consistently selected nearly identical tissue regions regardless of the chosen cluster number (**Supplementary Fig. 19**). This stability demonstrates the robustness of the ROI selection algorithm to moderate variations in tissue segmentation granularity.

To investigate the impact of the number of histology clusters and ROI size on ROI selection, we conducted a series of experiments using the previously introduced gastric cancer dataset. In this analysis, we segmented the histology image with cluster numbers set to 6, 9, 12, 15, 18, and 21. For each segmentation result, we selected ROIs of three different sizes: 2 mm ×2 mm, 4 mm ×4 mm, and 6 mm ×6 mm. As shown in **Supplementary Fig. 20**, S2Omics demonstrates greater robustness when the number of histology clusters or the ROI size increases. Notably, even with the smallest 2 mm ×2 mm ROIs, S2Omics successfully identified ROIs when provided with detailed histology segmentation, highlighting its effectiveness in selecting meaningful tissue regions.

### Computational efficiency of s2Omics

To effectively guide spatial omics experiments, an ROI selection algorithm must be computationally efficient, allowing users to receive feedback quickly. S2Omics extracts histology image features using foundation models such as UNI. However, since S2Omics captures local image features at near single-cell resolution, processing large H&E images, which often contain hundreds of millions of pixels, can be computationally intensive. To enhance computational efficiency, S2Omics employs a down sampling strategy, where superpixels are down sampled based on a predefined down sampling rate. Our systematic evaluation of S2Omics’s computational performance across various H&E image dimensions and down sampling rates (**Fig. 6c**) showed that a 1:100 down sampling rate provides an optimal balance between ROI quality and processing speed. With this approach, S2Omics can select ROIs for a 20 mm × 20 mm tissue section in just 15 minutes on an Nvidia Tesla V100 16GB Tensor Core GPU, significantly improving usability without compromising accuracy.

## Discussion

In this paper, we presented S2Omics, an automated framework that uses only H&E images to guide spatial omics experimental design. This tool addresses two critical challenges in current spatial molecular profiling studies: subjective ROI selection and inconsistent experimental design. By leveraging recently developed pathology image foundation models to extract histological features, S2Omics identifies the most representative regions for analysis. When prior knowledge is available, it can prioritize or de-emphasize specific tissue areas, increasing its adaptability to diverse experimental needs.

Beyond ROI selection, S2Omics provides a virtual preview of whole-tissue molecular organization through cell type and cell community label broadcasting. By extrapolating information from limited spatial molecular data within the selected ROIs, S2Omics reconstructs tissue-wide cellular architecture, offering users an early, tissue-scale perspective prior to full experimental profiling. This approach not only informs downstream experimental design but also addresses technical challenges such as discontinuous sampling in certain platforms such as CosMx. Together, the speed, flexibility, and tissue-wide inference capabilities make S2Omics a powerful tool for standardizing and optimizing spatial molecular profiling experiments.

While the current implementation is based on H&E images, S2Omics can be readily extended to incorporate other histological stains such as trichrome, Elastica van Gieson stain, or other stained images, for ROI selection. Furthermore, S2Omics can be adapted for ROI selection based on tissue regions defined by other molecular data. For instance, it can select ROIs based on cell types identified from PhenoCycler-Fusion and subsequently conduct spatial transcriptomics using the ROIs recommended by S2Omics^13^.

Although S2Omics has demonstrated strong performance, several limitations remain. First, it currently cannot accept DAPI or immunofluorescence images as input due to the lack of appropriate feature extraction models for these image types. Second, S2Omics relies on pathology image foundation models for feature extraction, which, while effective, require downsampling to maintain reasonable processing speed. Eliminating this need through more efficient feature extraction will be a priority for future development. Third, the current implementation handles only a small number of samples by concatenating them into a single composite image, a strategy that is not scalable for large-scale studies. Addressing these constraints will be essential for enabling S2Omics to fully support spatial omics design.

Given the growing interest in spatial omics, we anticipate that S2Omics will serve as a foundational tool for researchers, which enables them to allocate resources effectively by focusing on the most critical tissue regions and maximizing information gained from their experiments. This standardization is especially crucial for clinical and disease-focused studies, where consistent and reproducible outcomes are essential for generating meaningful and translatable biological insights. By addressing this fundamental yet overlooked step, S2Omics establishes a new standard for experimental rigor and reproducibility in spatial omics research.

## Data availability

We analyzed the following datasets: (1) 10x Genomics human kidney cancer and kidney non-diseased Xenium data (https://www.10xgenomics.com/datasets/human-kidney-preview-data-xenium-human-multi-tissue-and-cancer-panel-1-standard); (2) 10x Genomics human liver cancer and liver non-diseased Xenium data (https://www.10xgenomics.com/datasets/human-liver-data-xenium-human-multi-tissue-and-cancer-panel-1-standard); (3) 10x Genomics human breast cancer Xenium data, in situ sample 1 replicate 1, in situ sample 2 (https://www.10xgenomics.com/products/xenium-in-situ/preview-dataset-human-breast); (4) 10x Genomics human colorectal cancer and colon non-diseased Visium HD data, P1 CRC, P2 CRC, P3 NAT, P5 CRC (https://www.10xgenomics.com/products/visium-hd-spatial-gene-expression/dataset-human-crc); (5) human gastric cancer Xenium data generated by Tae Hyun Hwang lab (https://zenodo.org/records/15164980, https://upenn.box.com/s/nazcbustyepbf7dsv0ih8enxw3wr04×6); (6) human kidney CosMx data generated by Katalin Susztak lab (https://upenn.box.com/s/5zjynedjrachn1p8c4y1us1vonhgqm3d). (7) human breast cancer data generated by Anderson et al.^33^ (https://zenodo.org/records/3957257, G1, G2, and G3). (8) H&E image of breast cancer biopsies generated by Anupma Nayak lab (https://upenn.box.com/s/nrahq680fbr3aegm5uz3vz9yxyhspv4l). (9) H&E image of gastric cancer sample with signet-ring cells generated by Linghua Wang lab (https://upenn.box.com/s/psdx9qj6gmeiyr4kgedsdb9gvssbvcn4). Details of the datasets analyzed in this paper are described in **Supplementary Table 1**.

## Code availability

The S2Omics algorithm was implemented in Python and is available on GitHub at https://github.com/ddb-qiwang/S2Omics.

## Acknowledgments

M.L. was partly supported by the following NIH grants R01HG013185, R01LM014592, U19NS135582, R01HL171595, and U01CA294518. T.H.H. was partly supported by NIH grants R01CA276690 and U01CA294518, DOD grant CA190578, the Eric and Wendy Schmidt Foundation’s AI Innovation Award through the Mayo Clinic Foundation, and the Torrey Coast Foundation. L.W. was supported in part by the NIH National Cancer Institute grants U01CA294518, U01CA264583, R01CA266280, U24CA274274, and Break Through Cancer. L.W. is a member of the James P. Allison Institute and the Institute for Data Science in Oncology at The University of Texas MD Anderson Cancer Center and receives research funding from both institutes.

## Author contributions

This study was conceived of and led by M.L. M.Y. designed the model and algorithm with input from M.L., implemented the software, and led data analyses. K.J., H.Y., A.S., C.L., S.Y. T.H. performed data analyses. B.D., J.L., and K.S. provided the CosMx kidney data and helped with analyses and interpretation. J.R.C., I.J., M.K., J.H.P., T.H.H. provided the gastric cancer Xenium data and helped with analyses and interpretation. Y.L., I.L., L.M.S.S., and L.W. provided the gastric cancer Visium data, and I.L. and L.M.S.S. annotated this sample. A.N. provided and annotated the breast cancer biopsy samples. J.H.P. annotated the gastric cancer and colorectal cancer data. M.D., E.E.F., and P.W. provided feedback on the model and biological interpretations. M.Y. and M.L. wrote the paper with feedback from other co-authors.

## Competing financial interests

M.L. receives research funding from Biogen Inc. unrelated to the current manuscript. M.L. is a co-founder of OmicPath AI LLC. T.H.H. is a co-founder of Kure.ai therapeutics, and has received consulting fees from IQVIA; these affiliations and financial compensations are unrelated to the current manuscript. L.W. serves as a member of the Scientific Advisory Board for SELLAS Life Sciences and receives compensation outside the scope of this submitted work. The other authors declare no competing financial interests.

## Methods

S2omics consists of three components: a histology image feature extractor, a region of interest (ROI) selector, and a whole-slide cell type/cell community broadcaster. Below, we provide a detailed description of each component.

### Histology feature extractor

To facilitate the processing of histology images with different resolutions, we first rescale each image so that the size of one pixel is 0.5 µm × 0.5 µm. This rescaling ensures that each 16 × 16-pixel tile, or superpixel, corresponds to 8 µm × 8 µm, approximately the size of a single cell. To ease the subsequent tiling procedure, we pad the rescaled image so that its height and width are both divisible by 224. A simple quality control process is applied to all superpixels. Specifically, superpixels with high average RGB values and low RGB variance are filtered out as they typically correspond to background regions or areas without nuclei. While any histology image feature extraction method can be used to obtain features for each superpixel, in this paper, we chose to use UNI^21^.

Let *X* ∈ ℝ^*M*^ × ℝ^*N*^ × ℝ^3^ be the histology image with height *M* and width *N*, and three RGB color channels. We partition *X* into a (*M*/16)-row, *N*/16-column rectangular grid of 16 × 16-pixel image tiles: 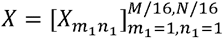, where each,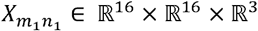. Next, we crop all 224 × 224-pixel neighborhood tiles that have a 16 × 16-pixel tile 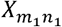 at its center: 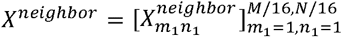, where each 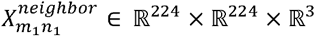. Centered on the small tile 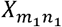, the large tile 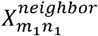, which encompasses about 200 cells, captures the tissue microenvironment surrounding the center cell. Extracting image features for all 16 × 16-pixel tiles and their corresponding 224 × 224-pixel neighborhood tiles can be time consuming. To reduce computational time, a down-sampling procedure can be applied to both *X* and,*X*^*neighbor*^ avoiding extracting histology image features for every superpixel. In practice, we recommend using a down sampling rate of 1/100 for ROI selection to effectively reduce computational cost. We show in **Fig. 6c** that this down-sampling strategy significantly reduces computational time while maintaining high accuracy in ROI selection.

In our recent iStar paper^34^, we demonstrated that hierarchically extracted image features are crucial for capturing both global and local image characteristics, which are essential for super-resolution gene expression prediction. However, foundation models like UNI lack the capability to hierarchically extract such image features. To mimic the hierarchical image feature extraction used in iStar, we employ a 224 × 224-to-16 × 16 vision transformer (ViT)^35^, denoted as *f* _*ViT*_. The ViT maps each 224 × 224-pixel neighborhood tile into a *C*_1_-dimensional feature vector,

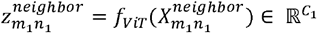

We introduce another function, 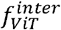, which extracts features of the 16 × 16-pixel tile located at the center of the current 224 × 224-pixel neighborhood tile. This function utilizes the output of the second-to-last layer of *f*_*ViT*_,yielding a *C*_2_ dimensional feature vector,

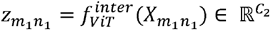

Next, we concatenate the local and neighborhood features to obtain a combined histology feature image 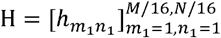 of *C*_1_ + *C*_2_ channels, where each 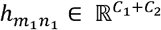 is the histology feature vector at superpixel (*m*_1_, *n*_1_).

In our implementation, we use UNI as *f*_*ViT*_, thus setting *C*_1_= *C*_2_ = 2048. User can replace UNI here with any pathology foundation model as long as its model structure is 224 × 224-to-16 × 16 ViT. In addition to the default UNI^36^, S2Omics also provides additional foundation models, including HIPT^22^, Prov-GigaPath^26^, and Virchow2^27^ as options. Our evaluations have shown that the computational efficiency varies substantially across models. HIPT-based feature extraction is ten times faster than UNI, due to its more compact architecture and smaller training dataset. In contrast, GigPath and Virchow2 require approximately three times longer processing times than UNI, consistent with their larger model sizes and computational complexity (**Supplementary Fig. 21**).

### ROI selector

The ROI selector is designed to identify regions of interest (ROIs) based on histology image segmentation. We consider two approaches for ROI selection: 1) without prior knowledge, where ROIs are chosen in an unbiased manner based solely on H&E image derived features, and 2) with prior knowledge, where biological or pathological information is incorporated to prioritize or de-emphasize specific tissue structures or cellular compositions during ROI selection.

#### Histology image segmentation

Once the histology image features are extracted, we segment the image into different clusters. First, we reduce the dimensionality of the histology image features using principal components analysis^37^. Each dimensionality reduced feature vector includes 80 principal components, representing histological information in a 16 × 16 superpixel tile. These superpixels serve as individual samples, which we then cluster using the K-means algorithm^38^. To capture hierarchical relationships among clusters and minimize the influence of outliers, we apply a two-stage hierarchical strategy. In the first stage, histological features are partitioned into a relatively large number of preliminary clusters using the K-means algorithm. A subsequent merging step consolidates over-segmented clusters based on image feature similarity. This approach minimizes the influence of outliers, which are later merged into stable and biologically meaningful regions in the final segmentation output. By default, S2Omics initializes with 20 clusters and merges them to about 15 under this procedure, a configuration that performed robustly across datasets in this study. We recommend this default and suggest adjusting parameters as needed for specific data types. Since histological patterns are strongly correlated with molecular variations, we will use these histology-based clusters to select ROIs. Spatial omics experiments conducted within these selected ROIs are expected to capture most molecular variations across the entire tissue slice.

While K-means serves as the default clustering algorithm due to its robust performance (**Supplementary Fig. 22**), S2Omics also supports agglomerative clustering, BIRCH, Bisecting K-means, Fuzzy C-means, Leiden, and Louvain as additional options, providing flexibility to accommodate diverse real-world datasets.

#### ROI selection without prior knowledge

We first illustrate our ROI selection algorithm when no prior knowledge about the tissue slice is available. In this case, the entire tissue slice is subject to ROI selection. S2Omics requires users to specify only the size of the ROI, after which the algorithm either identifies the optimal single ROI or automatically determines the optimal number and locations of multiple ROIs. For simplicity, we assume all ROIs are square-shaped and of the same size. However, the algorithm can be easily modified to accommodate ROIs of varying sizes and non-square shapes if needed. Hereafter, a ROI refers to a square shaped region with pre-specified size by users. Our goal is to select ROI(s) from the entire tissue slice so that the tissue variations, as represented by histological clusters, in the entire tissue can be captured by the selected ROI(s).

Suppose our goal is to select *R* optimal ROIs {ROI_1_,ROI_2_,…,ROI_*R*_ }, where we need a metric to evaluate how effectively these square-shaped regions capture the tissue variation in the whole-slide image. For simplicity, we call superpixels that passed quality control as valid superpixels in the following context. Ideally, selected ROIs should be representative of the tissue, meaning they should effectively capture tissue variations as reflected by histological clusters in the H&E image. To achieve this, the selected ROIs should satisfy two key criteria:

1. Maximizing tissue content – The selected ROIs should minimize empty regions and capture as much tissue as possible.
2. Ensuring cluster representativeness – The ROIs should include a diverse range of histological clusters, ensuring that both major and rare clusters are well represented. When no prior knowledge about the tissue is available, the ROIs should aim for equal representation of all tissue structures, capturing both common and rare histological patterns.

Guided by these principles, an ROI scoring metric should incorporate the following components:

1. The first component is the coverage score, *S*_coverage_, which quantifies the proportion of valid superpixels within the selected ROIs. It is defined as

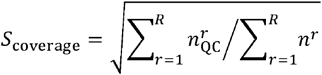

where *n*^*r*^ is the number of superpixels in the *r*-th ROI ROI_*r*_, and 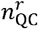 is the number of superpixels that passed quality control within that ROI. This score encourages the selection ROIs with densely distributed cells rather than cell-sparse regions. The square root transformation is applied to ensure sparsely populated histological clusters are not over penalized. This adjustment prevents the ROI selection strategy from systematically ignoring regions with lower cell densities, which may still hold important biological significance.
2. The second component is the balance score, *S*_balance_, which evaluates how well the histological cluster distribution in the selected ROIs aligns with an ideal uniform the distribution. Let *C*_ROIs_ = [*C*_1_,…, *C_K_*] represent the proportion of each histological cluster in the selected ROIs, and 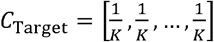 denote the ideal scenario where all clusters are equally represented. The balance score is defined as the cosine similarity between these two vectors:

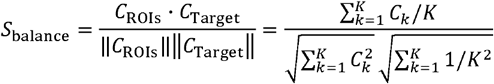

This score ensures that the selected ROIs capture a diverse range of histological clusters, with rare clusters receiving appropriate representation.

To create an effective ROI selection metric, we must also account for diminishing returns when selecting additional ROIs. As the number of ROIs increases, the marginal benefit of adding another ROI decreases when the experimental cost continues to rise. To address this, we introduce a size score, *S*_size_, which reflects the impact of the total ROI size on overall quality. The size score is defined as the inverse of a logit function over the effective sampling rate of valid superpixels within the selected ROIs:

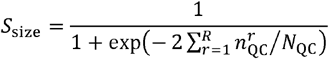

where 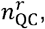 as mentioned previously, is the number of superpixels that passed quality control within that ROI, and *N*_*QC*_ is the total number of superpixels that passed quality control in the entire tissue slice. This function ensures that the score increases more slowly as the number of selected ROIs grows, balancing coverage and representativeness against experimental feasibility.

The final ROI score, *S*_ROI_, is defined as the weighted geometric mean of three component scores, including balance, coverage, and size (**Supplementary Fig. 23**):

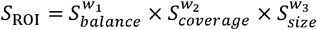

where *w*_1_, *w*_2_, *w*_3_ are tunable weights of non-negative real numbers that sum to 1, reflecting the relative importance of each component. In this study, we used the default setting 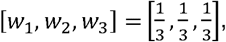 which prevents any single component from dominating the final score and ensures a balanced tradeoff between maximizing tissue content, preserving histological diversity, and controlling experimental cost. Users may adjust these weights to emphasize specific priorities for their experimental design.

#### ROI selection using ROI scores

Next, we describe the procedure for selecting ROIs using this metric. When *R* = 1, we begin by randomly sampling *L* candidate ROIs from all possible ROIs centered within the tissue section. The ROI with the highest *S*_ROI_ score is selected as the optimal ROI. We set *L* = 500 ×,,*A*_WSI_⁄*A*_ROI_, where,*A*_WSI_ and,*A*_ROI_ denote the areas of the whole tissue section and a single ROI, respectively. The rationale behind this choice is that when the ROI size is large or the overall tissue section is small, fewer randomly sampled ROIs are needed to ensure comprehensive coverage of the tissue section.

When R > 1, we need to address two questions: 1) determining the optimal number of ROIs, and 2) identifying the optimal locations for these ROIs. There are two natural approaches to solving this dual problem. One approach is to select all ROIs simultaneously, but this requires an extremely large number of samples, as the search space grows exponentially with *R*. The alternative approach is to selected ROIs sequentially, choosing ROI_r+1_ based on the previously selected ROIs {ROI_1_,ROI_2_,…,ROI*r*} However, this greedy strategy only guarantees local optimality at each step, which may lead to a suboptimal final set {ROI_1_,ROI_2_,…,ROI_R_ }. To balance computational efficiency with selection quality, we adopt a hybrid strategy:

- For *R* = 2, we select the best ROI pair {ROI_1_,ROI_2_ }simultaneously among from *L*^*2*^ randomly sampled pairs, where *L* is defined as before.
- For *R* = 2*m*, we divide the selection procedure into steps, *m* selecting the best ROI pair in each step from *L*^*2*^ randomly sampled pairs, conditioning on the previously selected pairs.
- For *R* = 2*m* + 1, we first determine the best {ROI_1_,ROI_2_,…,ROI _2*m*_}based on the pair-based selection strategy, and then add one final ROI from *L* random samples to maximize *S*_ROI_.

This hybrid approach ensures a more computationally feasible process while maintaining high-quality ROI coverage and representativeness.

#### Determination of the optimal number of ROIs

The optimal number of ROIs is determined through an iterative process, where we start from R = 1 upward until the change in *S*_ROI_ falls below a predefined threshold *τ* Specifically, when reaching the optimal ROI score *R*_optimal_, we ensure that:

- The increase in *S*_ROI_ from *R*_optimal_to *R*_optimal_ + 1 is smaller than *τ*.
- The increase in *S*_ROI_ from *R*_optimal_ to *R*_optimal_ 2 is smaller than 2 *τ*.

For all ROI selection experiments, we set *τ*.at 0.03, while for FOV selection experiments, *τ*.was set to 0 to enforce a stricter selection criterion.

#### ROI selection with prior knowledge

Experienced pathologists can refine ROI selection by leveraging prior knowledge. To mimic this capability, S2Omics offers an option for users to incorporate prior information into the ROI selection process. Specifically, after histology clusters are identified, users can designate certain clusters to be emphasized or discarded. The new balance score, 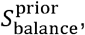 is calculated using the modified target proportion vector, 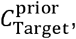 which incorporates prior information. Suppose that among the *K* histology clusters, the user intends to emphasize clusters *i*_1_,*i*_2_, …,*i*_*p*_ while excluding clusters *j*_1_,*j*_2_, …,*j*_*q*_ from the ROI section. In this case, the prior target cluster proportion vector is defined as:

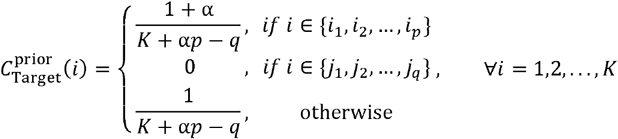

Here 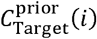 denotes the target proportion of cluster *i*, and α is the prior preference parameter that quantifies the user’s desired degree of emphasis. Specifically, clusters designated as positive priors are treated as α times more important than standard clusters. This formulation can be readily generalized to cases involving both positive and negative priors across multiple histological clusters. The final ROI score, *S*_ROi_, is then calculated using this revised balance score, while the overall ROI selection strategy remains consistent with the approach used in the absence of prior knowledge.

#### ROI selection for multiple tissue sections

S2Omics also supports ROI selection across multiple tissue sections. When multiple H&E images are available, S2Omics standardizes them by padding each image to uniform dimensions and concatenating them horizontally to construct a composite mega-image. This preprocessing step transforms multi-section data into a single-section format, allowing the same feature extraction and ROI selection procedures used for individual sections to be applied consistently. In doing so, S2Omics ensures a unified and reproducible analytical workflow regardless of the number of sections under study.

### Whole-slide cell type/cell community predictor

Once the ROIs are selected, spatial omics experiments will be conducted within these regions. The resulting data will enable us to characterize cell types present in the ROIs, and this information can subsequently be used to broadcast cell types and cell communities across the remaining portions of the tissue that were not subjected to spatial omics measurements. The whole-slide information predictor contains two autoencoders, one for cell type label broadcasting and the other for cell community label broadcasting. A cell community is defined as a group of cells with similar cell type compositions within their surrounding neighborhoods. Specifically, for each cell, its neighborhood is determined by the 500 nearest neighboring cells, identified using Euclidean distance based on their spatial locations. Similar concepts have been proposed by Schurch et al.^39^ and others^40^, which have demonstrated that cell communities often exhibit stronger association with clinically relevant phenotypes compared to traditionally obtained cell clusters. Below we describe the process for predicting cell type labels from histology images.

Suppose we have *n* cells within the selected ROIs, where each cell belongs to one of *C*_y_ different cell types. These cells’ cell type labels are given by *y* = {*y*_1_,*y*_2_, …,*y*_*n*_}and the corresponding histological features are *h* = {*h*_1_,*h*_2_, …,*h*_*n*_} To reduce the dimensionality of the histological features, we introduce an autoencoder consisting of an encoder,*f*_enc_, which maps the high-dimensonal histological features to a lower-dimensional latent space, and a decoder, *f*_dec_ that reconstructs the original histological features from the latent representations. Additionally, we use a predictor *f*_cls_ to classify the cell type labels based on the latent embeddings from the autoencoder. The training objective consists of two components, including a reconstruction loss

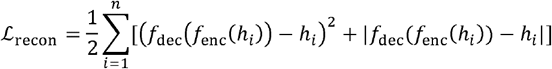

and a generalized cross-entropy loss which is widely utilized for training neural networks with noisy labels^41^,

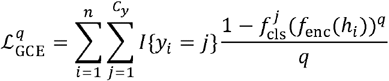

Here 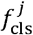 represents the predicted probability for cell type *j* and *q* is a parameter controlling the generalized cross entropy loss 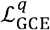 and was fixed at 0.6 in all experiments. The total loss function is given by

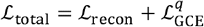

This ensures that the model learns a meaningful representation of histological features while accurately predicting cell types. Once this prediction model is trained, we can predict the cell type labels of the whole slide using the histology image features 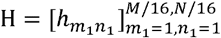 as input.

Cell community labels can be predicted using a similar approach.

### Evaluation criteria for ROI selection quality

We assess the quality of ROI selection using the following metrics: ROI score, cell type broadcasting accuracy, and cell community broadcasting accuracy. For tissue sections with manually selected ROIs informed by pathologist expertise, we introduce an additional metric that quantifies the overlap percentage between these ROIs and our selections. For tissue sections with paired CosMx data, which S2Omics leverages to enhance cell type spatial distribution, we assess the consistency between the predicted cell type distribution using all CosMx data and the prediction based solely on CosMx data within the selected ROI/FOV(s). All metrics range from 0 to 1, with higher values indicating better performance.

### Handling of fragile tissue samples

Tissue handling and sectioning parameters, such as fixation protocol, tissue age, hydration state, and sectioning thickness, are context-specific and should be optimized in consultation with local histologists or other domain experts. For S2Omics analysis, the required H&E-stained whole-slide image can serve as a reference for evaluating sample fragility and the risk of detachment or fragmentation during downstream processing. For high-risk specimens, users may generate multiple candidate ROIs and compare their associated ROI scores. The final ROI can then be selected in coordination with domain experts to balance tissue preservation with experimental objectives.

