## Supplementary information for "Designing smart spatial omics experiments with S2Omics"

**Supplementary Table 1. Datasets analyzed in this paper.**

| Species | Tissue | Data source | H&E image physical size | Protocol |
|---|---|---|---|---|
| Human | Gastric cancer | Generated by Tae Hyun Hwang lab (https://zenodo.org/records/15164980) | 24 mm × 10 mm | Xenium |
| Human | Gastric cancer | Generated by Tae Hyun Hwang lab (https://upenn.box.com/s/nazcbustyepbf7dsv0ih8enxw3wr04x6) | 25 mm × 22 mm | H&E only |
| Human | Gastric cancer | Generated by Linghua Wang lab (https://upenn.box.com/s/psdx9qj6gmeiyr4kgedsdb9gvssbvcn4) | 10.8 mm × 20.6 mm | H&E only |
| Human | Kidney non-diseased | 10x Genomics (https://www.10xgenomics.com/datasets/human-kidney-preview-data-xenium-human-multi-tissue-and-cancer-panel-1-standard, non-diseased) | 9 mm × 3 mm | Xenium |
| Human | Kidney cancer | 10x Genomics (https://www.10xgenomics.com/datasets/human-kidney-preview-data-xenium-human-multi-tissue-and-cancer-panel-1-standard, kidney cancer) | 8 mm × 2 mm | Xenium |
| Human | Liver non-diseased | 10x Genomics (https://www.10xgenomics.com/datasets/human-liver-data-xenium-human-multi-tissue-and-cancer-panel-1-standard, non-diseased) | 12 mm × 8 mm | Xenium |
| Human | Liver cancer | 10x Genomics (https://www.10xgenomics.com/datasets/human-liver-data-xenium-human-multi-tissue-and-cancer-panel-1-standard, liver cancer) | 9 mm × 4 mm | Xenium |
| Human | Breast cancer | 10x Genomics (https://www.10xgenomics.com/products/xenium-in-situ/preview-dataset-human-breast, in situ sample 1 replicate 1) | 10 mm × 8 mm | Xenium |
| Human | Breast cancer | 10x Genomics (https://www.10xgenomics.com/products/xenium-in-situ/preview-dataset-human-breast, in situ sample 2) | 9 mm × 6 mm | Xenium |
| Human | Colorectal cancer | 10x Genomics (https://www.10xgenomics.com/products/visium-hd-spatial-gene-expression/dataset-human-crc, VisiumHD sample p1 CRC) | 20 mm × 16 mm | VisiumHD |
| Human | Colorectal cancer | 10x Genomics (https://www.10xgenomics.com/products/visium-hd-spatial-gene-expression/dataset-human-crc, Xenium In Situ sample p1 CRC) | 20 mm × 16 mm | Xenium |
| Human | Colorectal cancer | 10x Genomics (https://www.10xgenomics.com/products/visium-hd-spatial-gene-expression/dataset-human-crc, VisiumHD sample p2 CRC) | 21 mm × 13 mm | VisiumHD |

| | | | | |
|---|---|---|---|---|
| Human | Colon non-diseased | 10x Genomics (https://www.10xgenomics.com/products/visium-hd-spatial-gene-expression/dataset-human-crc, VisiumHD sample p3 NAT) | 22 mm × 12 mm | VisiumHD |
| Human | Colorectal cancer | 10x Genomics (https://www.10xgenomics.com/products/visium-hd-spatial-gene-expression/dataset-human-crc, VisiumHD sample p5 CRC) | 20 mm × 18 mm | VisiumHD |
| Human | Kidney (T2D and normal) | Generated by Katalin Susztak lab (https://upenn.box.com/s/5zjynedjrachn1p8c4y1us1vonhgqm3d) | 21 mm × 19 mm | CosMx |
| Human | Breast cancer | Published with paper 'Spatial deconvolution of HER2-positive breast cancer delineates tumor-associated cell type interactions' (https://zenodo.org/records/3957257, G1) | 9 mm × 8 mm | ST |
| Human | Breast cancer | Published with paper 'Spatial deconvolution of HER2-positive breast cancer delineates tumor-associated cell type interactions' (https://zenodo.org/records/3957257, G2) | 9 mm × 8 mm | ST |
| Human | Breast cancer | Published with paper 'Spatial deconvolution of HER2-positive breast cancer delineates tumor-associated cell type interactions' (https://zenodo.org/records/3957257, G3) | 9 mm × 8 mm | ST |
| Human | Breast cancer | Generated by Anupma Nayak lab (https://upenn.box.com/s/nrahq680fbr3aegm5uz3vz9yxyhspv4l) | 24 mm × 22 mm | H&E only |

**Supplementary Fig. 1 | Application to a gastric cancer tissue section. a,** Heatmap referring to the percentage of cell types in each histology cluster. **b,** Heatmap of cell type enrichment in cell communities.

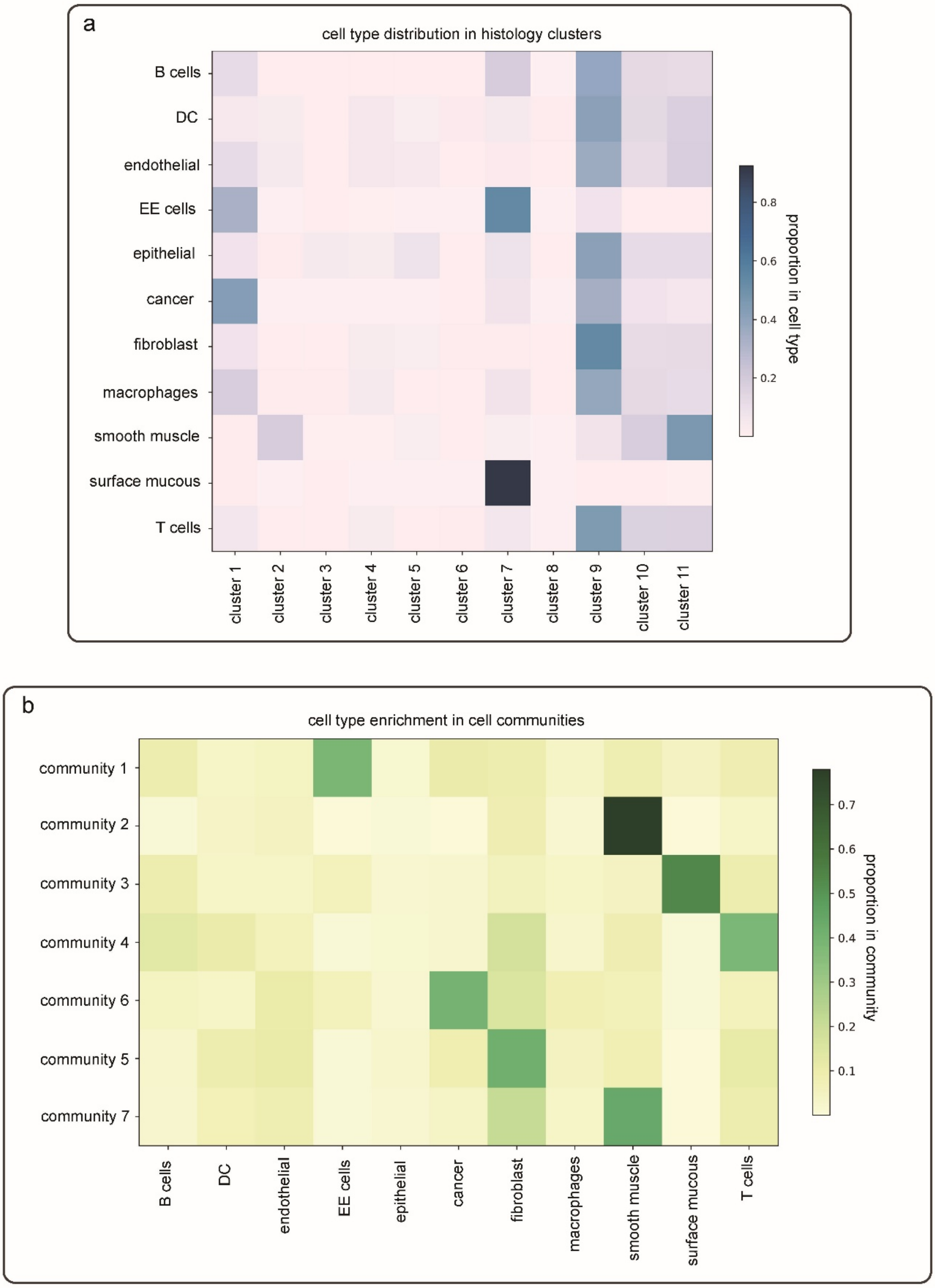

**Supplementary Fig. 2 | Application to a gastric cancer tissue section. a,** True positive (TP) and false positive (TP) superpixels of S2Omics's TLS prediction and cancer cell prediction using the cell type and cell community annotations within the selected ROI. **b,** Sankey diagrams between Xenium-based cell type/cell community annotation (left) and S2Omics's prediction from the selected ROI (right). **c,** Quantitative evaluation of S2Omics's ROI selection against 500 randomly sampled ROIs within the tissue section, covering the full range of all possible selection.

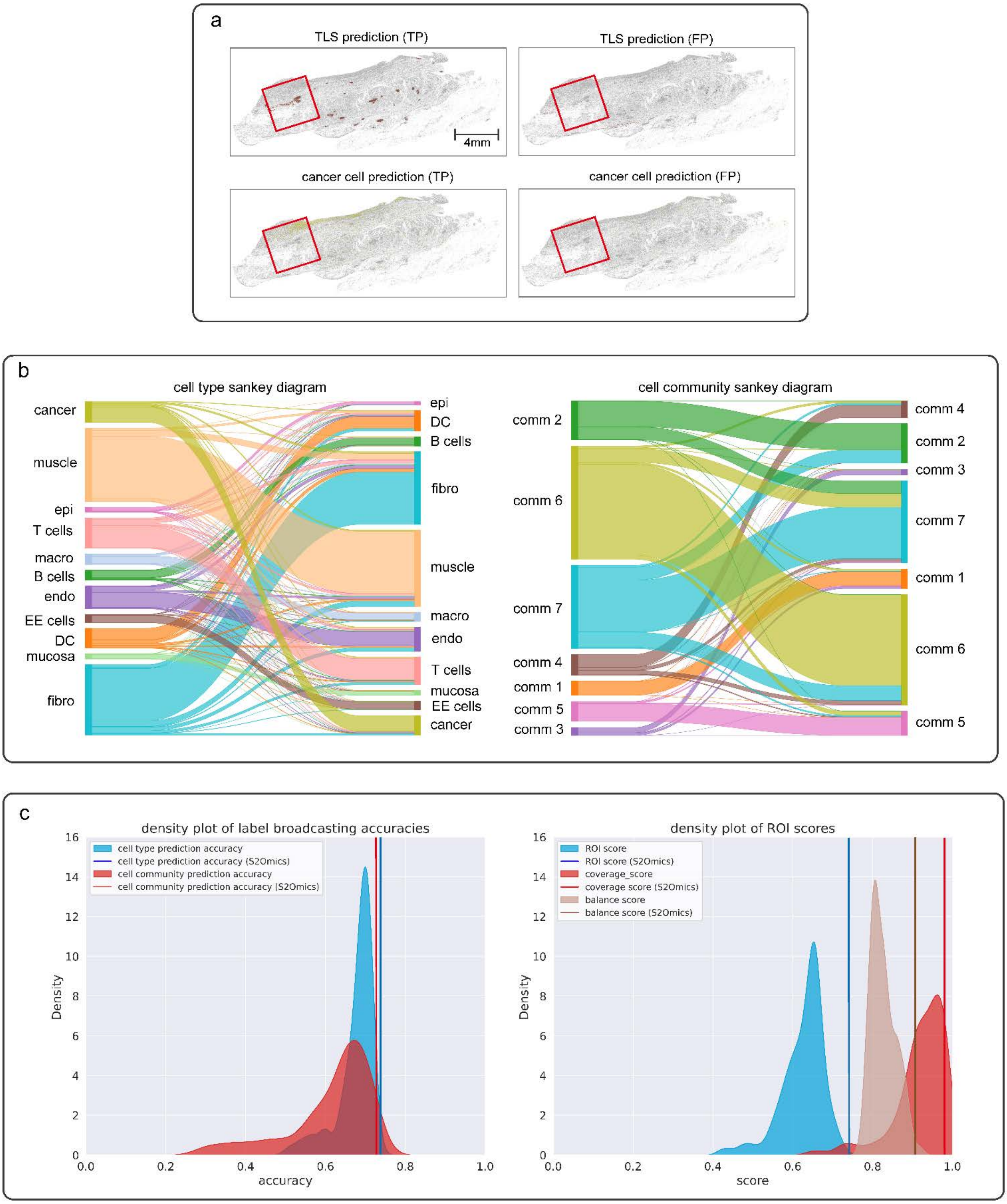

**Supplementary Fig. 3 | Example of a less favorable ROI in a gastric cancer tissue section. a,** Visualization of a less favorable ROI, the histology cluster and cell community compositions of the ROI. **b,** Cell type and cell community prediction from the ROI. For all superpixels that have Xenium-based cell type annotation, corresponding histology features and annotations were served as training data. Cell type labels of all superpixels that passed quality control were predicted using the trained model. **c,** Sankey diagrams between Xenium-based cell type/cell community annotation (left) and S2Omics's prediction from the selected ROI (right).

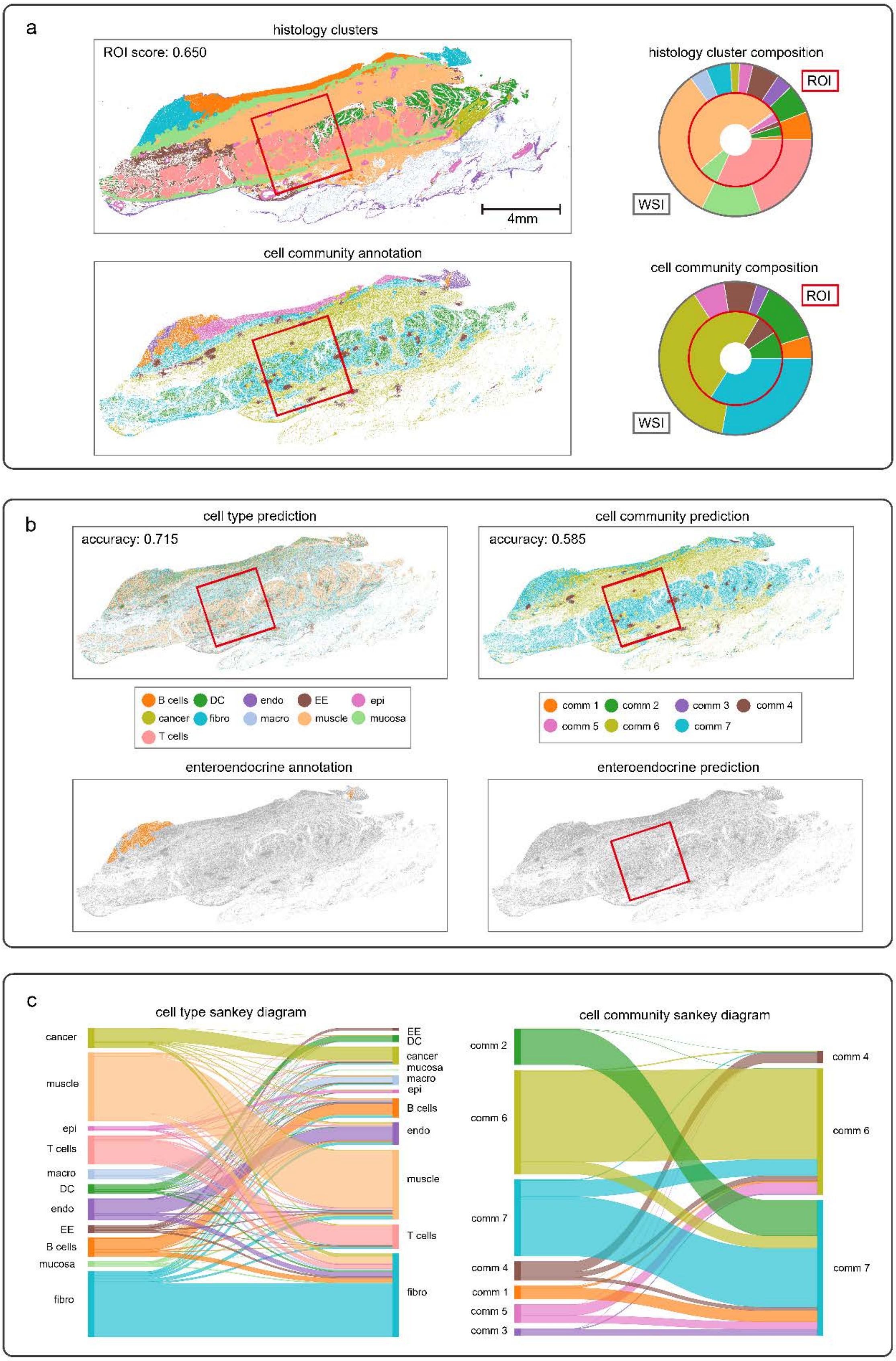

**Supplementary Fig. 4 | Application to a colorectal cancer tissue section. a,** Visual comparison between ROIs separately selected by S2Omics (red box) and experienced pathologist (dashed red box) on a colorectal cancer tissue section (P2 CRC). **b,** Histology cluster compositions of the two ROIs and their pie chart visualizations. **c,** Visium HD-based cell type annotation and S2Omics's cell type prediction on the whole tissue section. For all superpixels that have Visium HD-based cell type annotation, their histology features and annotations were served as training data. Cell type labels of all superpixels that passed quality control were predicted using the trained model. **d,** Distribution of cell types in histology clusters. Each square shows the percentage of cells in according cell type category and histology cluster occupied in all cells in that cell type category. **e.** the pathologist annotation and predicted cell type distribution of the 'ulceration with inflammation' region.

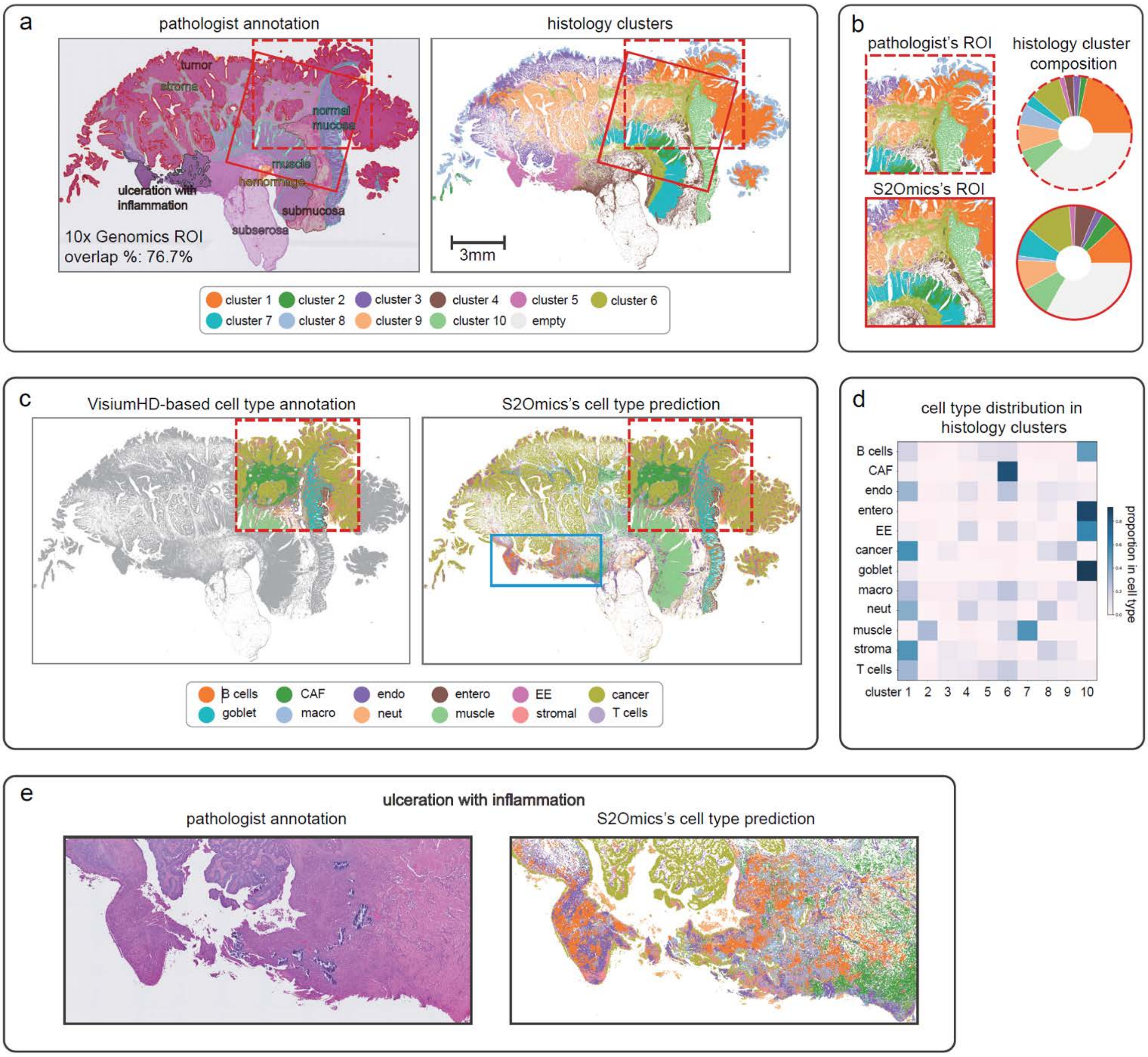

**Supplementary Fig. 5 | Application to a healthy colon tissue section. a,** Visual comparison between ROIs separately selected by S2Omics (red box) and experienced pathologist (dashed red box) on a healthy colon tissue section (P3 NAT). **b,** Histology cluster compositions of the two ROIs and their pie chart visualizations. **c,** Visium HD-based cell type annotation and S2Omics's cell type prediction on the whole tissue section. For all superpixels that have Visium HD-based cell type annotation, their histology features and annotations were served as training data. Cell type labels of all superpixels that passed quality control were predicted using the trained model. **d,** Distribution of cell types in histology clusters. Each square shows the percentage of cells in according cell type category and histology cluster occupied in all cells in that cell type category.

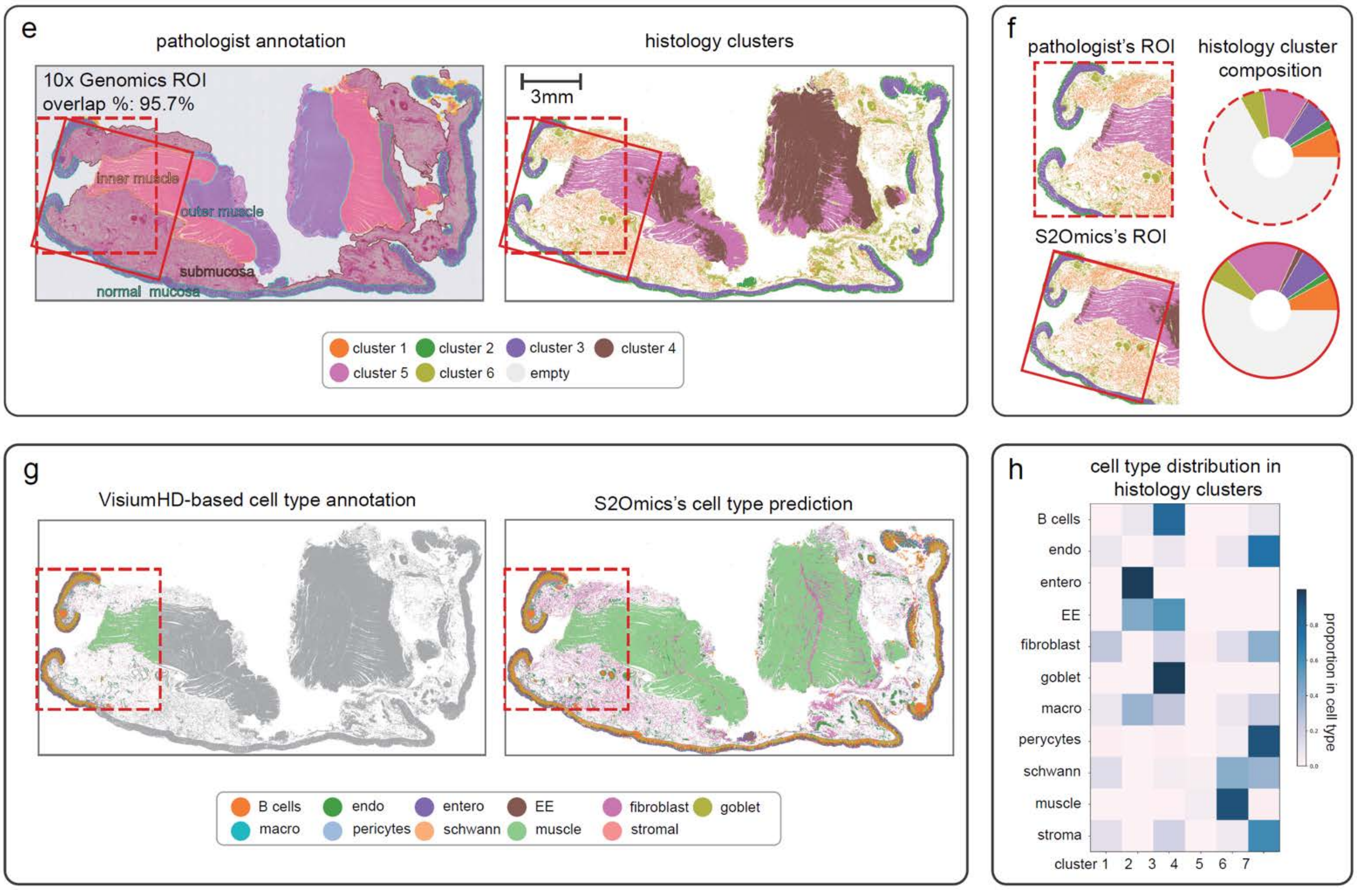

**Supplementary Fig. 6 | Application to a breast cancer tissue section. a,** H&E-stained image and histology segmentation obtained by S2Omics. 2 mm × 2 mm ROIs (red box) selected by S2Omics. **b,** Visual comparison between histology cluster and cell community compositions of WSI (outer ring) and S2Omics's ROI (inner ring). **c,** Cell type enrichment in cell communities. **d,** Distribution of cell types in histology clusters. Each square shows the percentage of cells in according cell type category and histology cluster occupied in all cells in that cell type category. **e,** Visual comparison between the ground truth and S2Omics-predicted cell labels from the selected ROI. For all superpixels that have Xenium-based cell type and cell community annotation, according histology features and annotations were served as training data for cell type and cell community predictors. Cell type and cell community labels of all superpixels that passed quality control were predicted using the trained models. **f,** Sankey diagrams between Xenium-based cell type/cell community annotation (left) and S2Omics's prediction from the selected ROI (right).

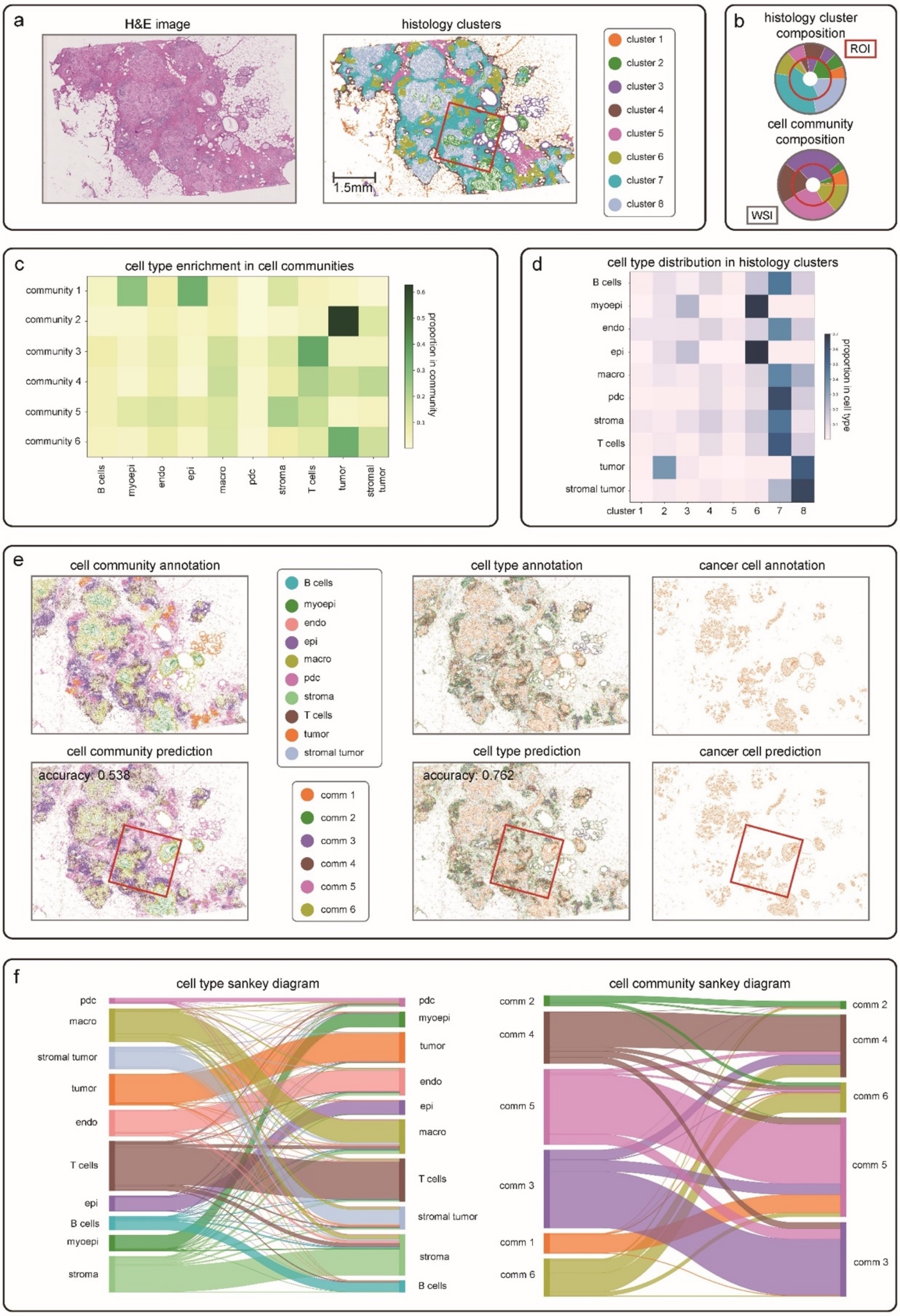

**Supplementary Fig. 7 | Application to a liver cancer tissue section. a,** H&E-stained image and histology segmentation obtained by S2Omics. 1.5 mm × 1.5 mm ROIs (red box) selected by S2Omics. **b,** Visual comparison between the ground truth and S2Omics-predicted cell labels from the selected ROI. For all superpixels that have Xenium-based cell type and cell community annotation, according histology features and annotations were served as training data for cell type and cell community predictors. Cell type and cell community labels of all superpixels that passed quality control were predicted using the trained models. **c,** Cell type enrichment in cell communities. Distribution of cell types in histology clusters. Each square shows the percentage of cells in according cell type category and histology cluster occupied in all cells in that cell type category. **d,** Visual comparison between histology cluster and cell community compositions of WSI (outer ring) and S2Omics's ROI (inner ring). **e,** Distribution of cancer cell in tissue section and S2Omics's prediction. f, Sankey diagrams between Xenium-based cell type/cell community annotation (left) and S2Omics's prediction from the selected ROI (right).

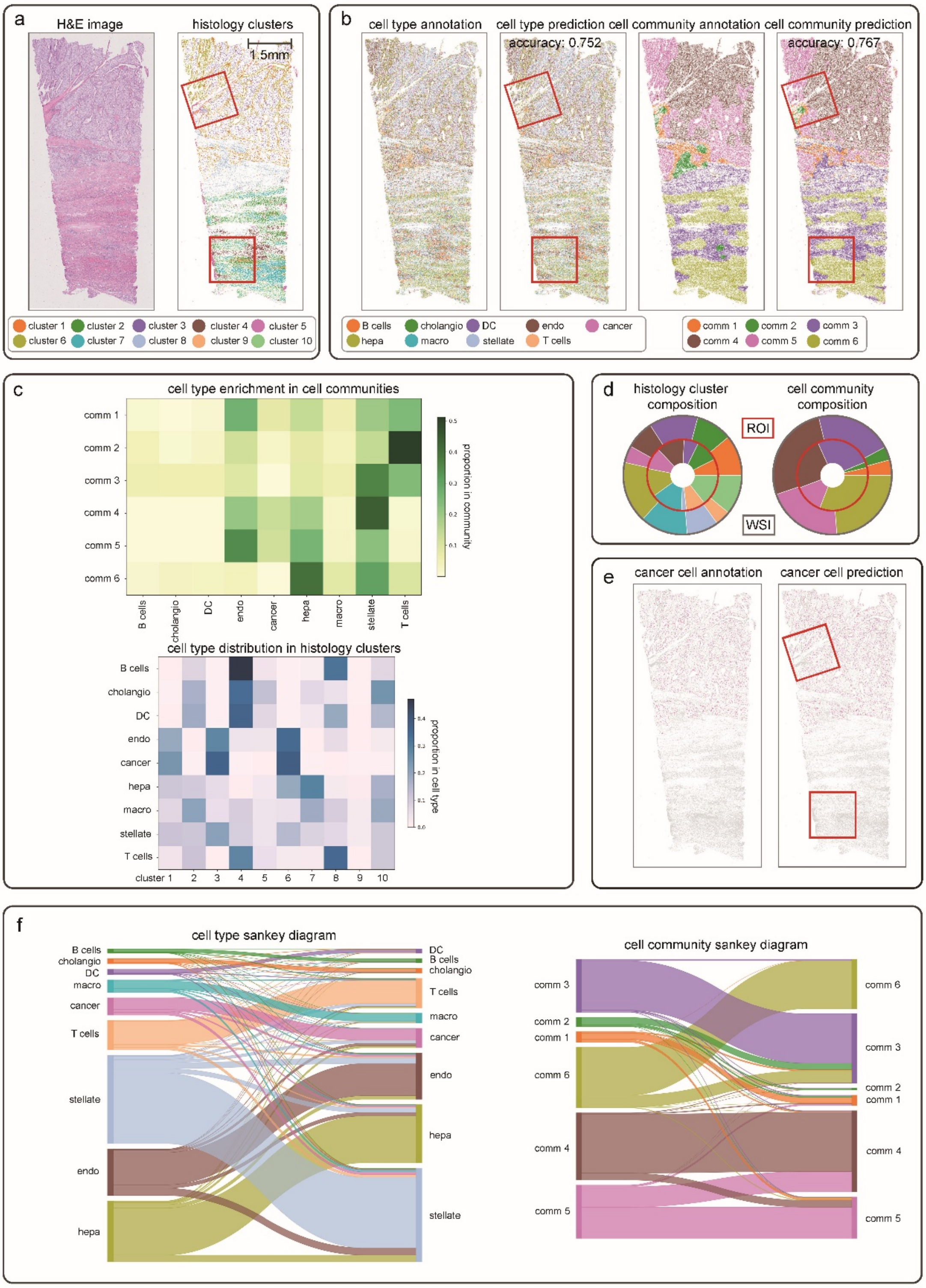

**Supplementary Fig. 8 | Application to a kidney cancer tissue section. a,** H&E-stained image and histology segmentation obtained by S2Omics. 1.5 mm × 1.5 mm ROIs (red box) selected by S2Omics. **b,** Visual comparison between the ground truth and S2Omics-predicted cell labels from the selected ROI. For all superpixels that have Xenium-based cell type and cell community annotation, according histology features and annotations were served as training data for cell type and cell community predictors. Cell type and cell community labels of all superpixels that passed quality control were predicted using the trained models. **c,** Cell type enrichment in cell communities. Distribution of cell types in histology clusters. Each square shows the percentage of cells in according cell type category and histology cluster occupied in all cells in that cell type category. **d,** Visual comparison between histology cluster and cell community compositions of WSI (outer ring) and S2Omics's ROI (inner ring). **e,** Distribution of cancer cell in tissue section and S2Omics's prediction. **f,** Sankey diagrams between Xenium-based cell type/cell community annotation (left) and S2Omics's prediction from the selected ROI (right).

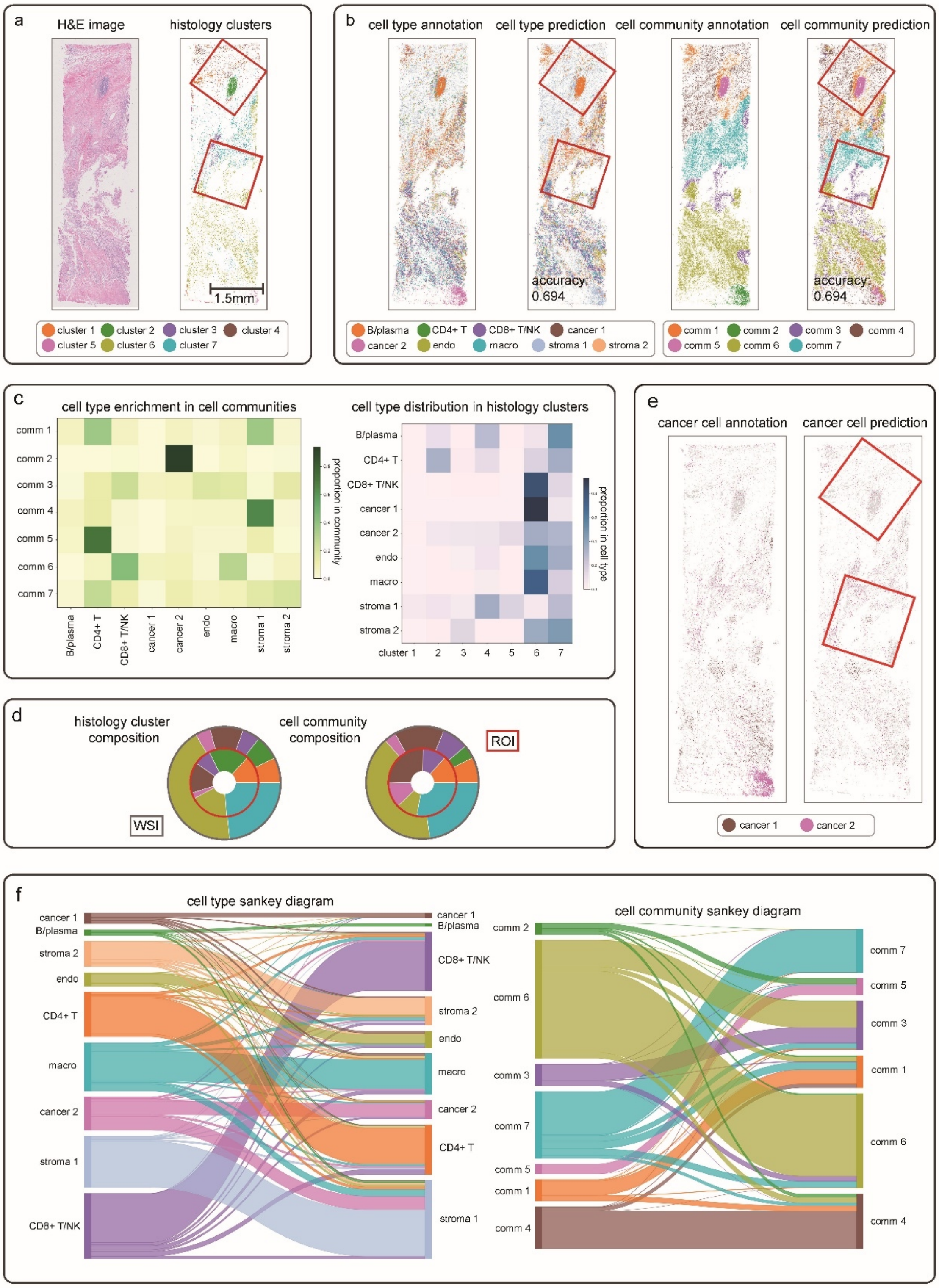

**Supplementary Fig. 9 | Application to a healthy kidney tissue section. a,** H&E-stained image and histology segmentation obtained by S2Omics. 1.5 mm × 1.5 mm ROIs (red box) selected by S2Omics. **b,** Cell type enrichment in cell communities. Distribution of cell types in histology clusters. Each square shows the percentage of cells in according cell type category and histology cluster occupied in all cells in that cell type category. **c,** Visual comparison between histology cluster and cell community compositions of WSI (outer ring) and S2Omics's ROI (inner ring). **d,** Visual comparison between the ground truth and S2Omics-predicted cell labels from the selected ROI. For all superpixels that have Xenium-based cell type and cell community annotation, according histology features and annotations were served as training data for cell type and cell community predictors. Cell type and cell community labels of all superpixels that passed quality control were predicted using the trained models. **e,** Sankey diagrams between Xenium-based cell type/cell community annotation (left) and S2Omics's prediction from the selected ROI (right).

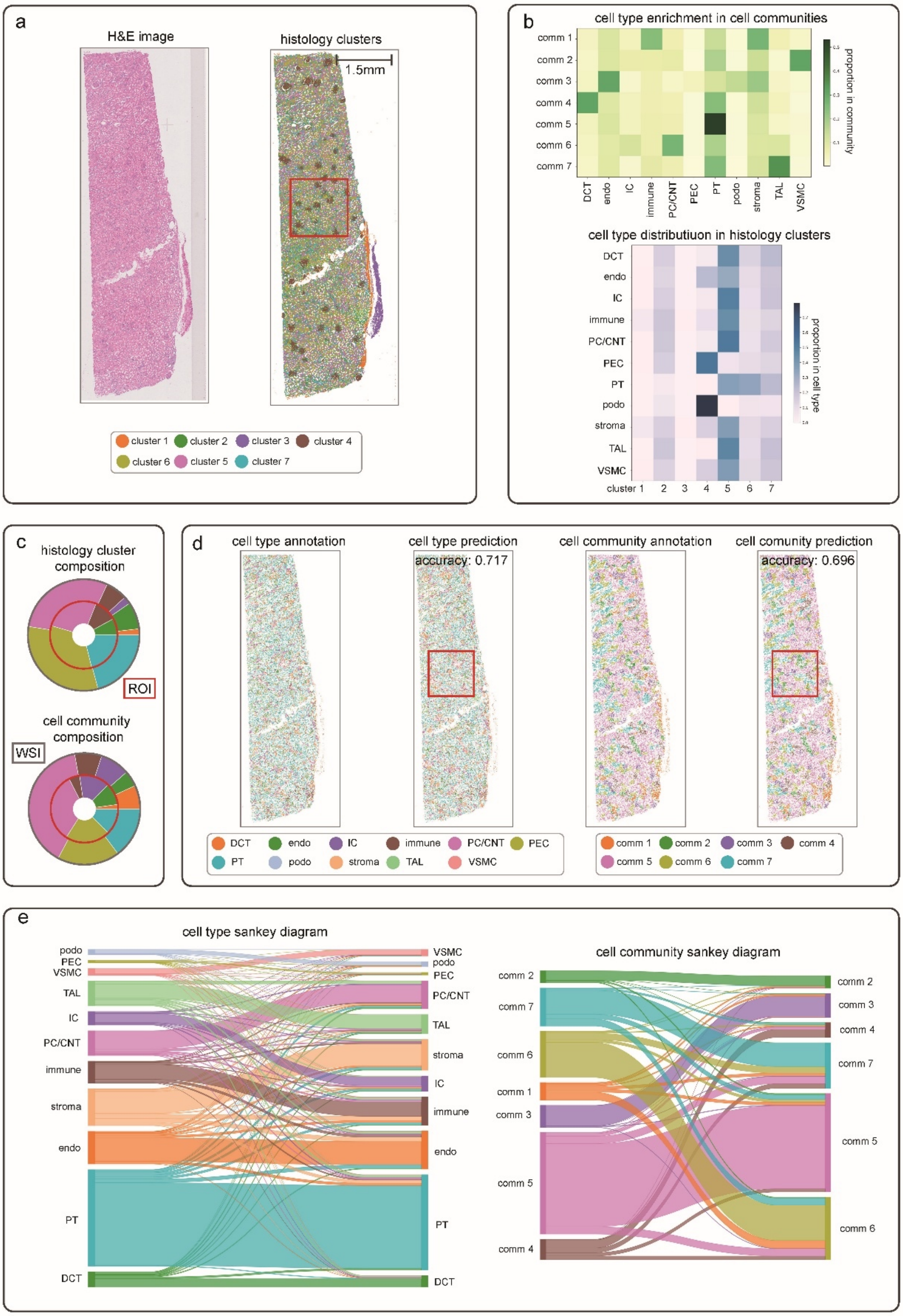

**Supplementary Fig. 10 | Application to a healthy liver tissue section. a,** H&E-stained image and histology segmentation obtained by S2Omics. 2 mm × 2 mm ROIs (red box) selected by S2Omics. **b,** Cell type enrichment in cell communities. Distribution of cell types in histology clusters. Each square shows the percentage of cells in according cell type category and histology cluster occupied in all cells in that cell type category. **c,** Visual comparison between histology cluster and cell community compositions of WSI (outer ring) and S2Omics's ROI (inner ring). **d,** Visual comparison between the ground truth and S2Omics-predicted cell labels from the selected ROI. For all superpixels that have Xenium-based cell type and cell community annotation, according histology features and annotations were served as training data for cell type and cell community predictors. Cell type and cell community labels of all superpixels that passed quality control were predicted using the trained models. **e,** Sankey diagrams between Xenium-based cell type/cell community annotation (left) and S2Omics's prediction from the selected ROI (right).

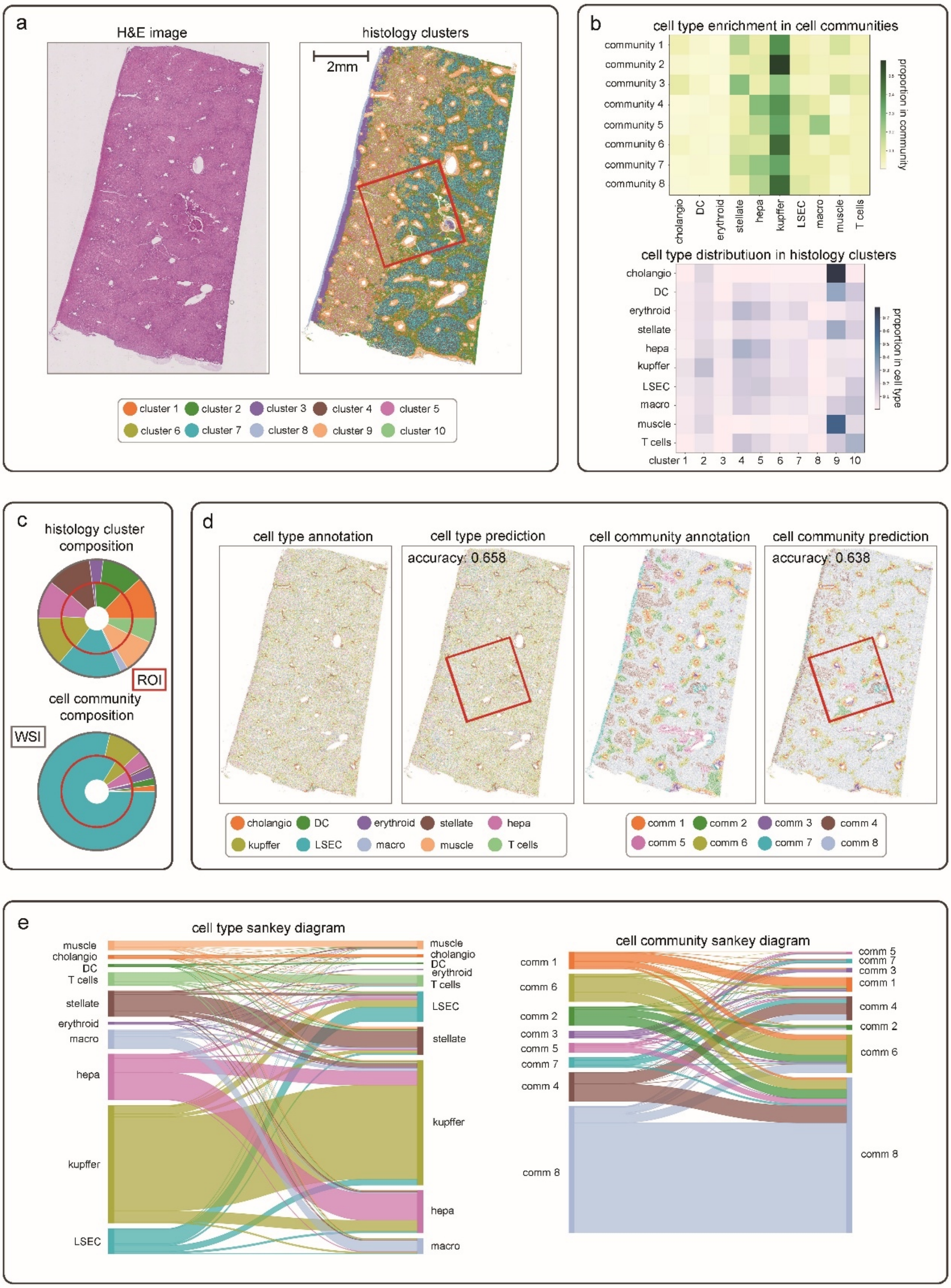

**Supplementary Fig. 11 | Application to a slide containing multiple breast cancer biopsies.** 1 mm × 1 mm ROIs selected by S2Omics visualized on pathologist annotation and histology clusters identified by S2Omics. The histology cluster composition of selected ROIs is similar to, but more balanced than the histology cluster composition of the whole slide.

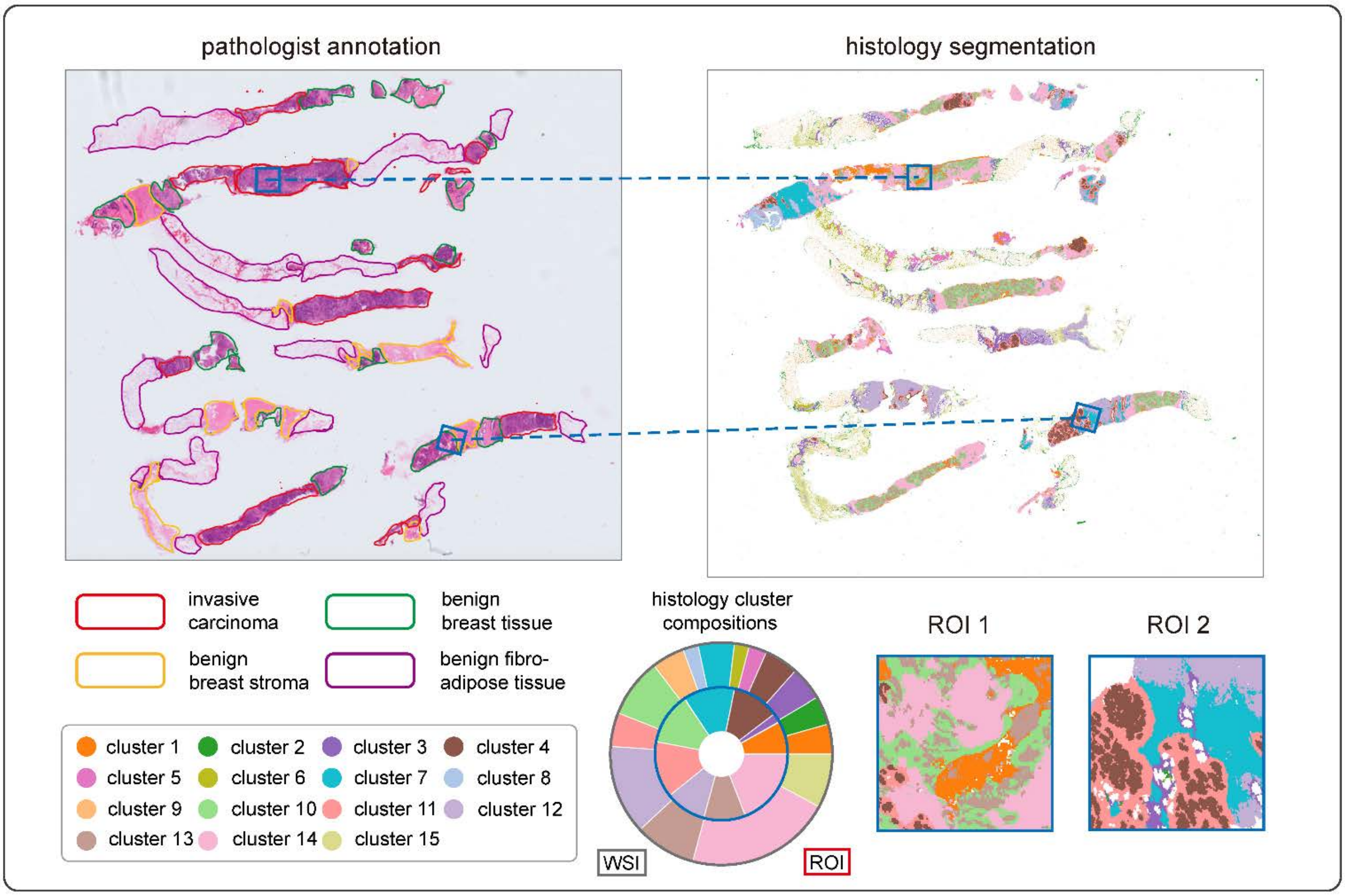

**Supplementary Fig. 12 | Application to a slide containing multiple breast cancer biopsies.** 1 mm × 1 mm ROIs selected by S2Omics visualized on pathologist annotation and histology clusters identified by S2Omics. Positive prior was given to the invasive carcinoma associated histology clusters (cluster 1, 10, 13, 14).

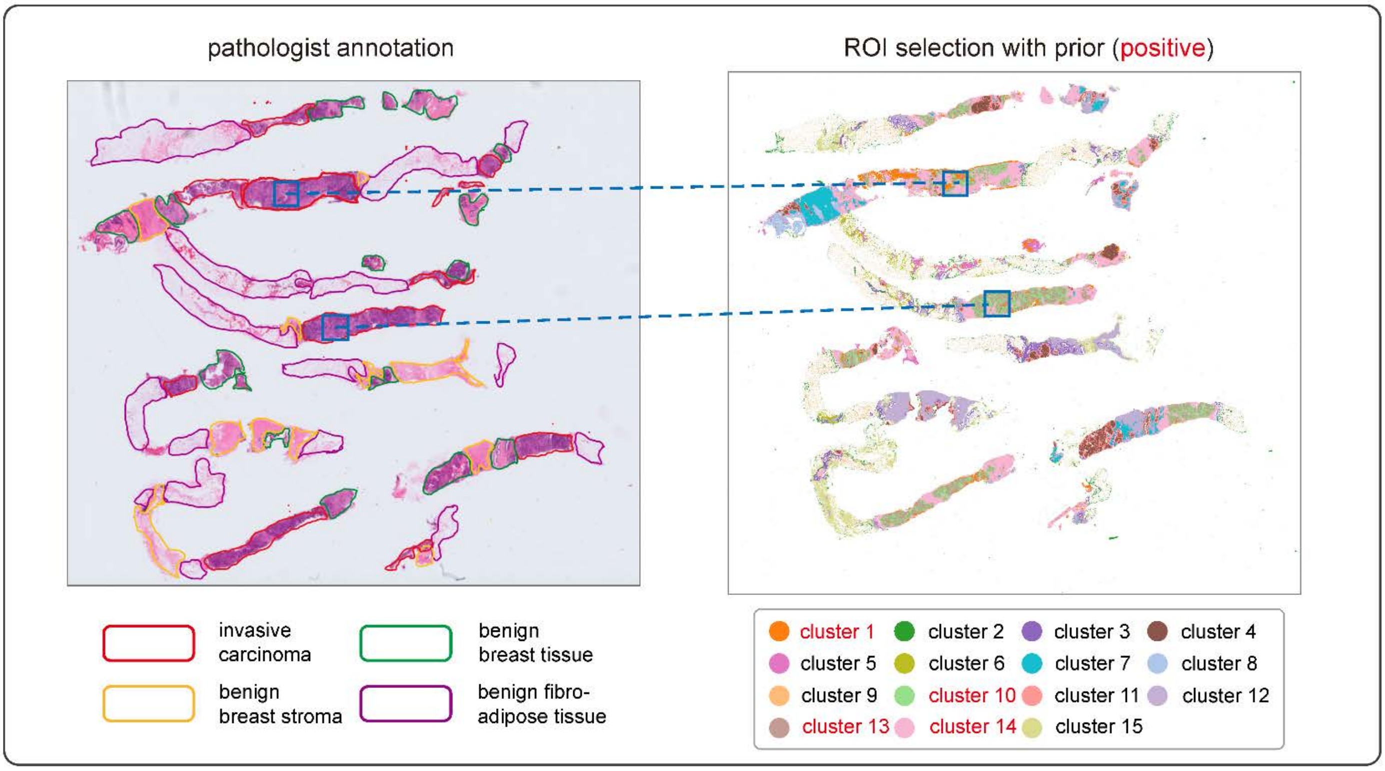

**Supplementary Fig. 13 | Selecting geometric ROIs on a slide containing multiple breast cancer biopsies.** 1mm × 2mm rectangular ROI and circular ROI with 1 mm diameter selected by S2Omics, visualized on the histology clusters identified by S2Omics. Positive prior was given to the invasive carcinoma associated histology clusters (clusters 1, 10, 13, 14).

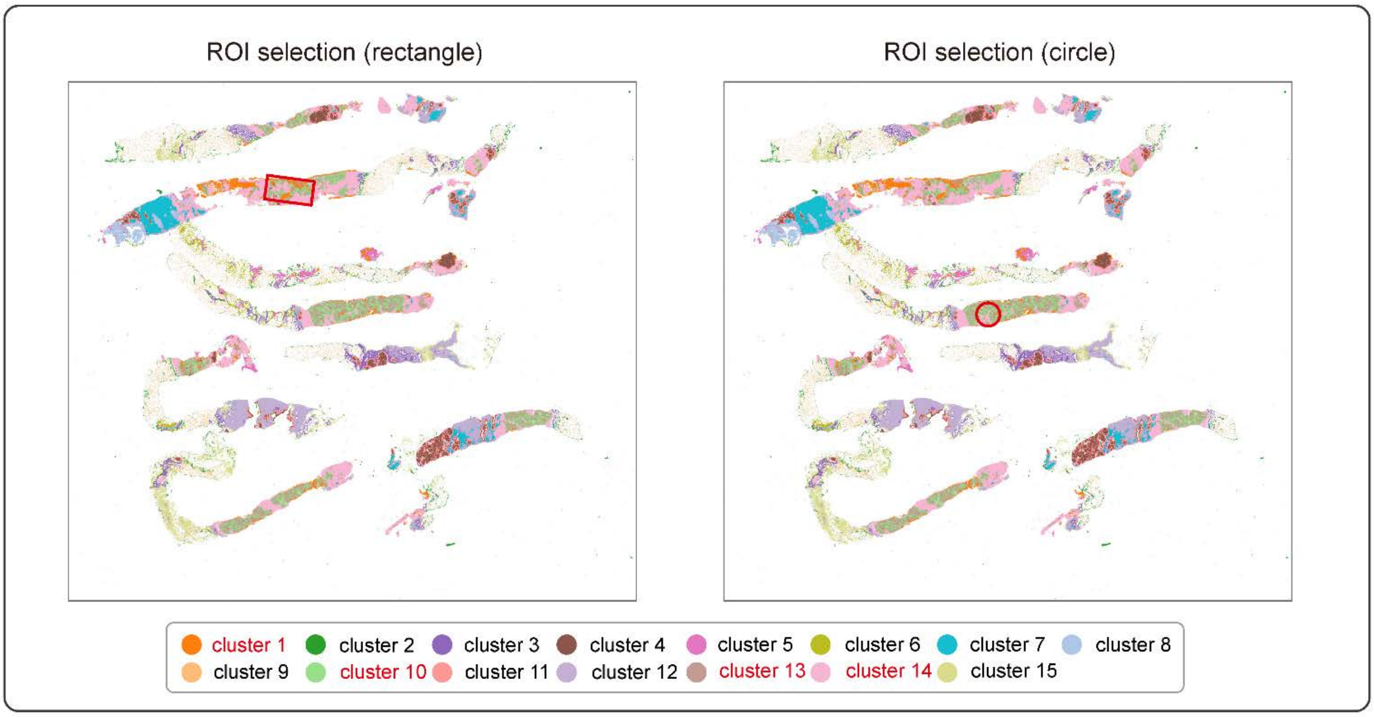

**Supplementary Fig. 14 | Application to a breast cancer data with consecutive tissue sections. a,** Pathologist annotation and H&E images of the consecutive sections (G1, G2, G3). **b,** 1.5 mm × 1.5 mm ROI selected by S2Omics, visualized on the joint histology segmentations of the consecutive sections obtained with S2Omics.

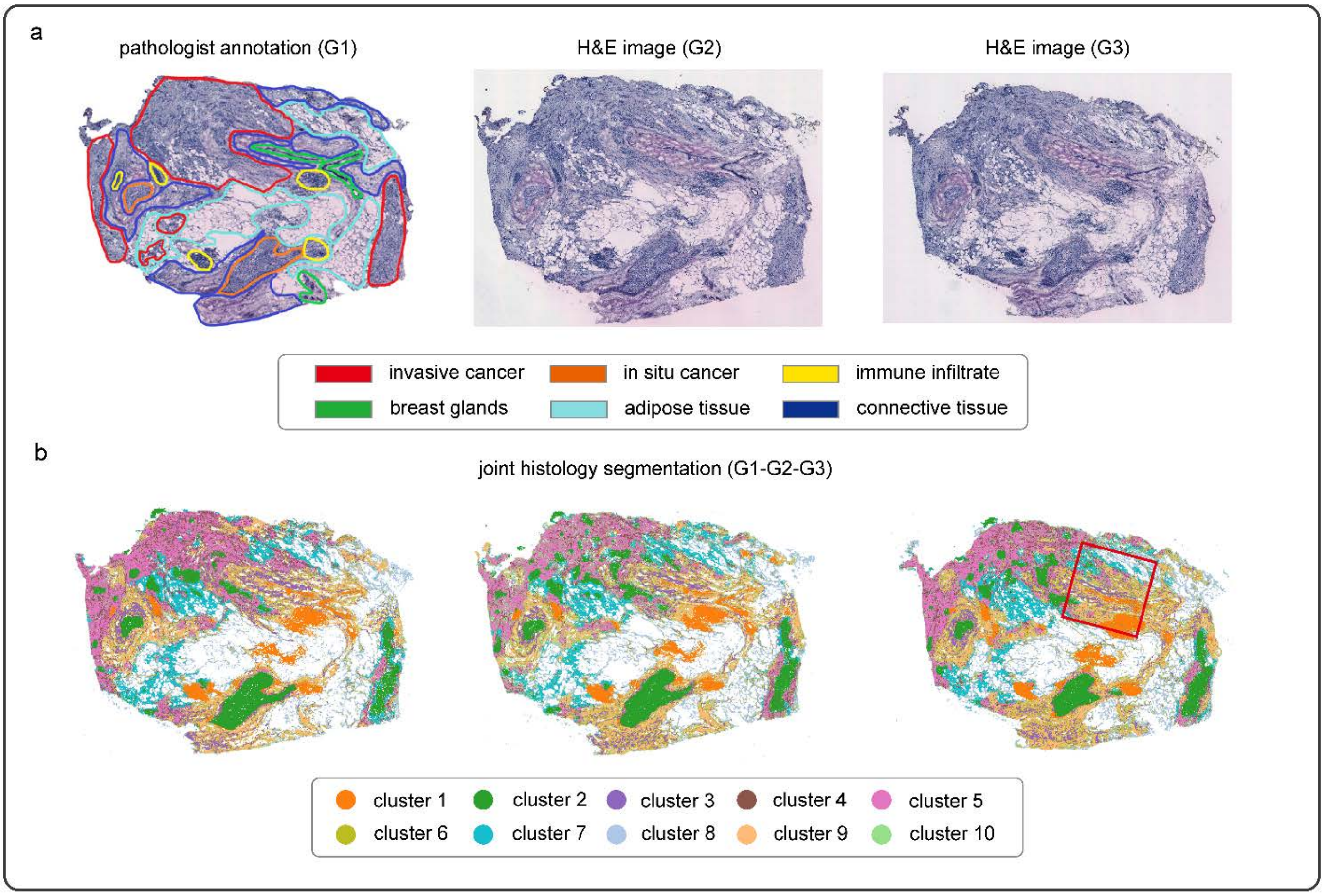

**Supplementary Fig. 15 | Application to a colorectal data with consecutive tissue sections. a,** H&E images of the consecutive sections (slice 1, slice 2). slice 2 stained after Xenium experiment. **b,** 6.5 mm × 6.5 mm ROI selected by S2Omics, visualized on the joint histology segmentations of the consecutive sections obtained with S2Omics. **c,** Cell type prediction maps for both tissue sections. The red dashed box in the left panel indicates the ROI manually selected by the 10x pathologist, within which the Visium HD experiment was performed. Cell type labels derived from Visium HD data within the pathologist-selected ROI were exclusively used for training the S2Omics label broadcasting module.

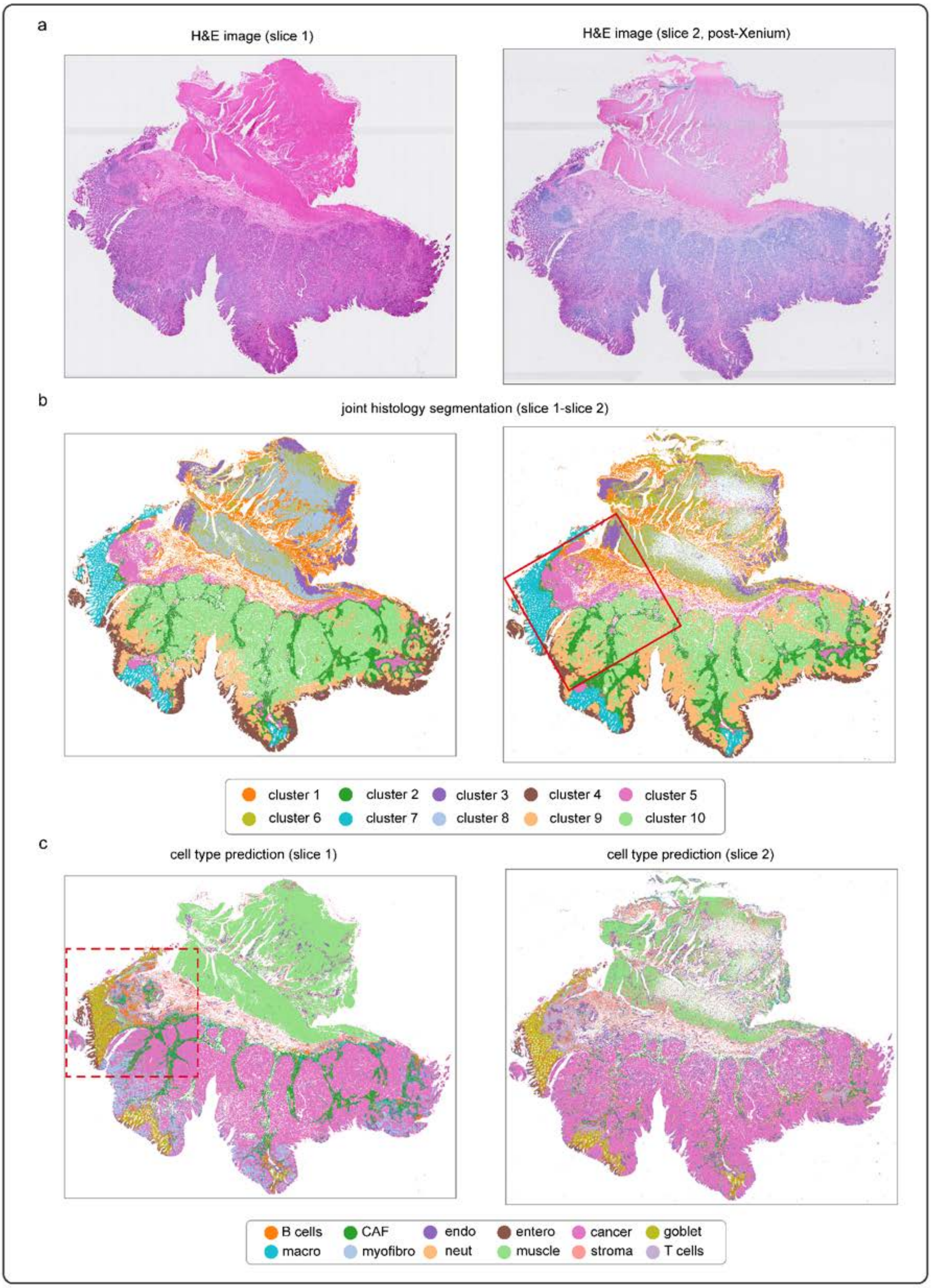

**Supplementary Fig. 16 | Application to a gastric cancer section containing a small proportion of signet-ring cells. a,** H&E image with magnified views of signet ring cells from three distinct regions within the tissue section. **b,** Pathologist annotation of each Visium spot based on the H&E image. GSE: gastric surface epithelium; GMG: gastric mucous glands. **c,** Histology clusters identified from the H&E image using S2Omics. **d,** Manually annotated signet-ring cells and histology clusters 8 and 10, showing significant spatial colocalization.

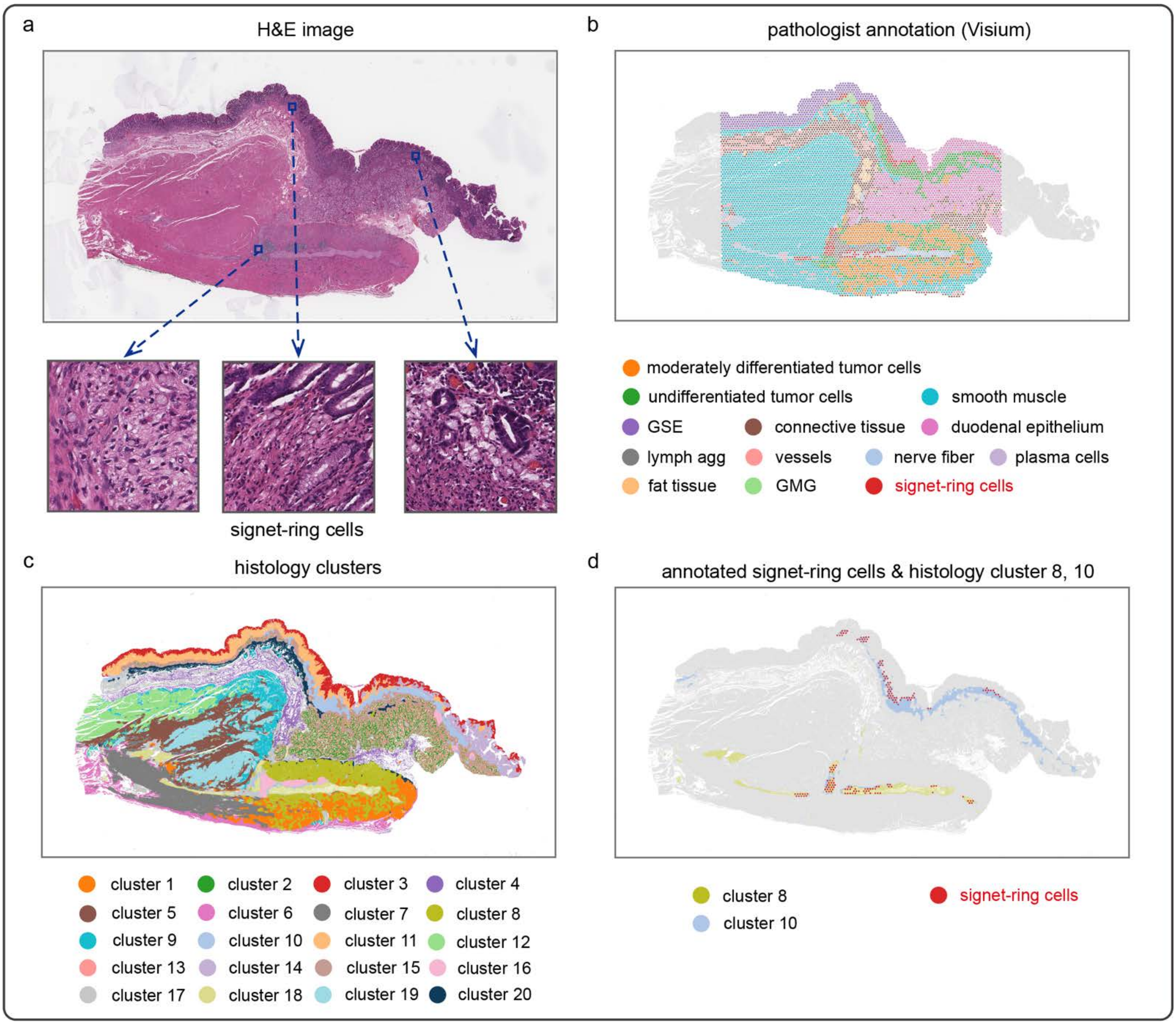

**Supplementary Fig. 17 | Application to a gastric cancer section containing a small proportion of signet-ring cells.** ROIs selected by S2Omics, visualized on H&E image, pathologist annotation of each Visium spot (GSE: gastric surface epithelium; GMG: gastric mucous glands), and the spatial distribution of signet-ring cells. The ROI size was set as 2 mm × 2 mm, 4 mm × 4 mm, and 6.5 mm × 6.5 mm, respectively.

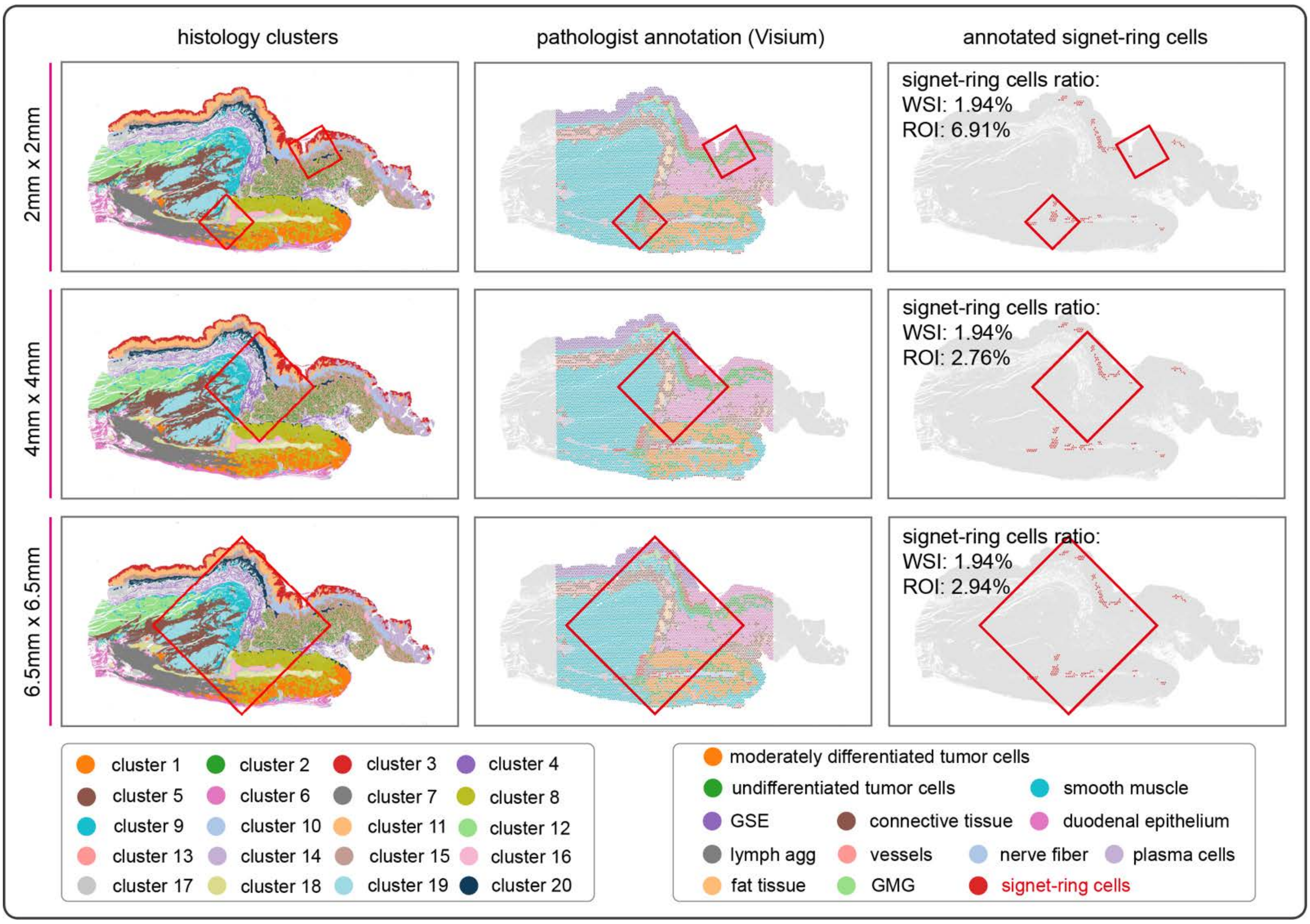

**Supplementary Fig. 18 | Application to a breast cancer section containing both CD4+ T and CD8+ T cells. a,** ROIs selected by S2Omics, visualized on the histology clusters identified from the H&E image using S2Omics. **b,** ROIs selected by S2Omics, visualized on the cell type annotation map of the tissue section derived from spatial transcriptomics (Xenium) data. **c,** histology clusters 4 and 7 are spatially highly correlated with T cells. **d,** ROIs selected by S2Omics, visualized on the spatial distribution and annotation CD4+ T and CD8+ T cells.

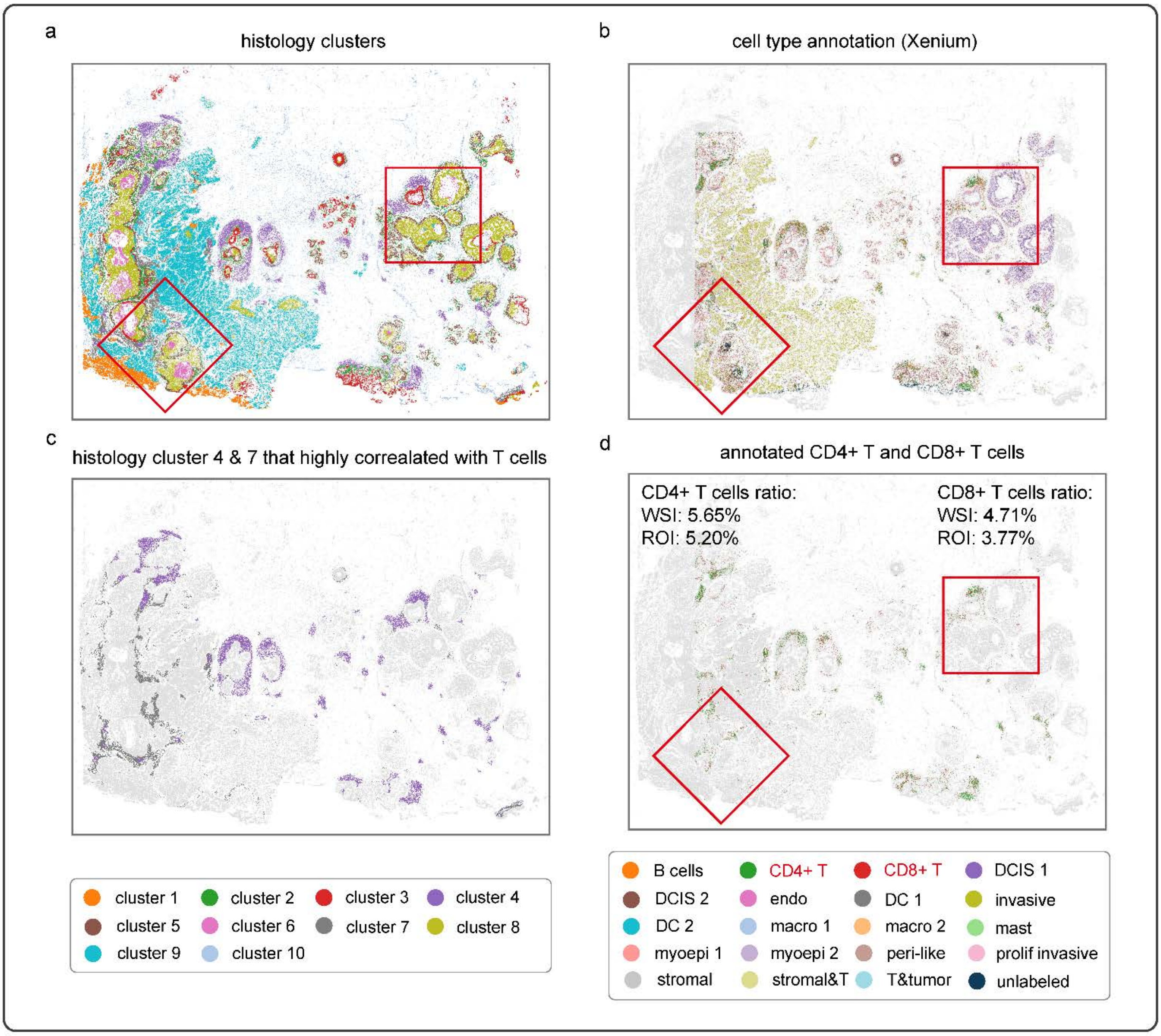

**Supplementary Fig. 19 | Impact of the number of final histology clusters on ROI selection in a gastric cancer section. Left,** H&E image with Euclidean distances between initial histology clusters and the merging thresholds. **Right,** ROIs selected by S2Omics, visualized on histology clusters obtained with different numbers of final clusters (after merging).

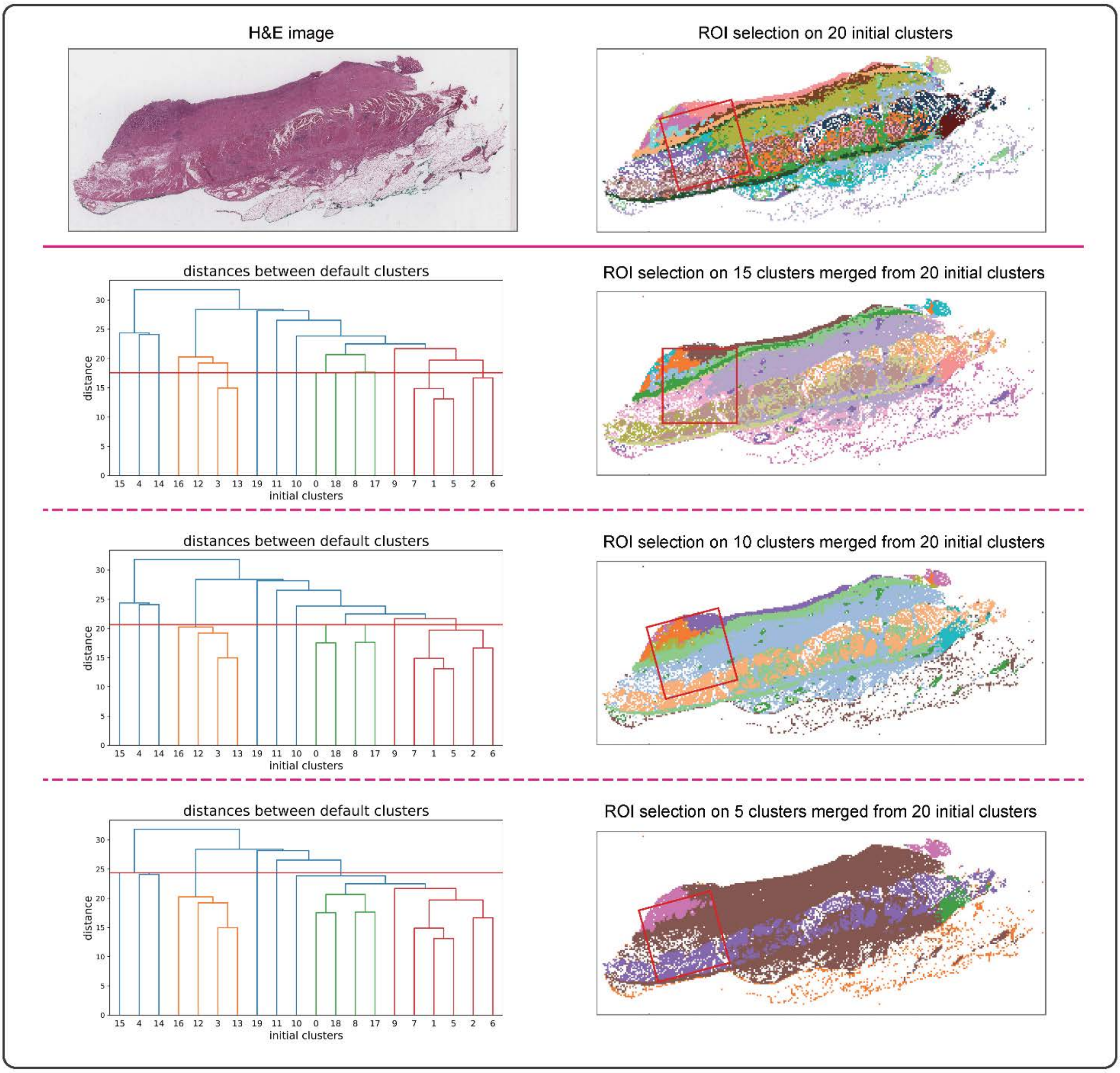

**Supplementary Fig. 20 | Evaluating the impact of histology cluster number and ROI size.** Histology segmentations and selected ROIs with varying cluster counts and ROI sizes. S2Omics was applied to a gastric cancer tissue section, segmenting it into 6, 9, 12, 15, 18, and 21 histology clusters. After obtaining the histology segmentations, S2Omics selected ROIs of sizes 2 mm × 2 mm, 4 mm × 4 mm, and 6 mm × 6 mm for each segmentation result.

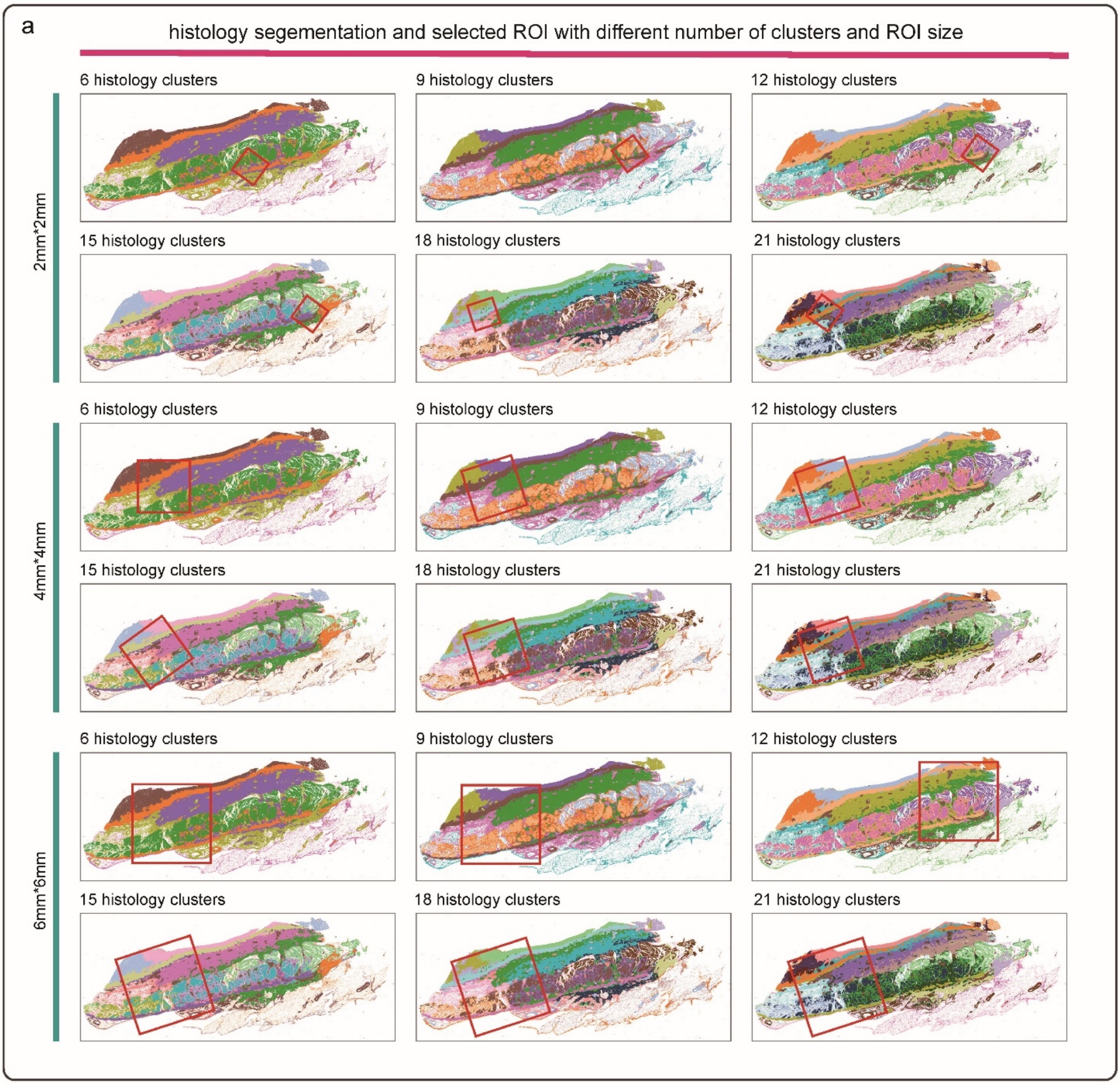

**Supplementary Fig. 21 | Comparative analysis of ROI selection and label broadcasting performance across S2Omics implementations using different foundation models for histological feature extraction. a,** Comparison between pathologist annotations and ROIs selected by S2Omics based on histological segmentations. Three different foundation models, including HIPT, Prov-GigaPath, and Virchow2, were employed for histological feature extraction. **b,** Cell type annotation maps derived from Xenium spatial transcriptomics data (ground truth) alongside cell type broadcasting results generated by different S2Omics implementations. For each S2Omics variant, the label broadcasting module was trained exclusively using cells within the ROI selected by that specific variant, along with their corresponding extracted histological features and cell type labels. Accuracy metrics were calculated using the Xenium-based cell type annotations as the reference standard.

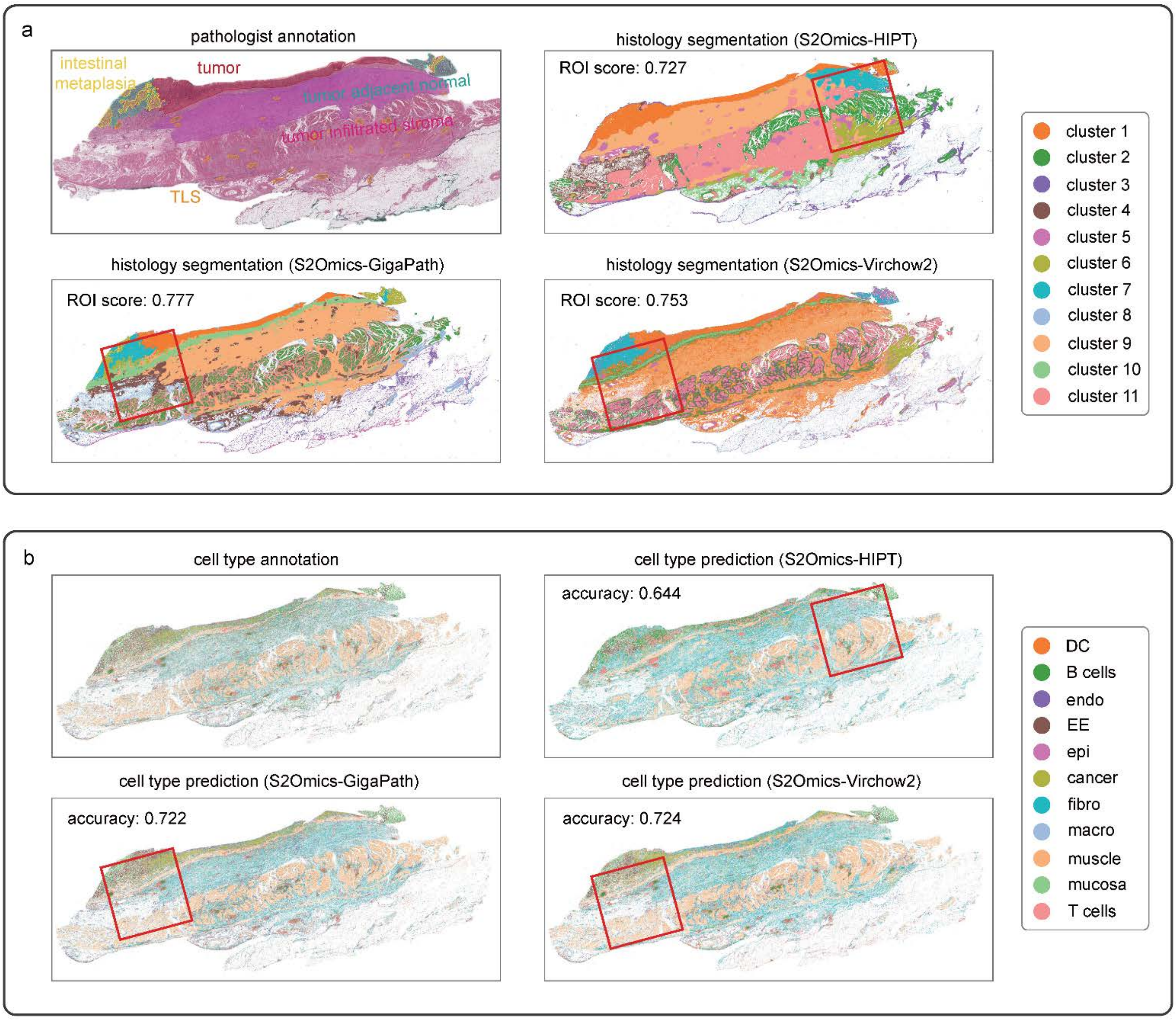

**Supplementary Fig. 22 | Impact of clustering methods on ROI selection in a colorectal cancer section.** **a,** ROIs selected by S2Omics using histology segmentations obtained by different clustering algorithms, visualized on the histology clusters. **b,** Line charts showing S2Omics performance on ROI selection and histology segmentation across different clustering algorithms.

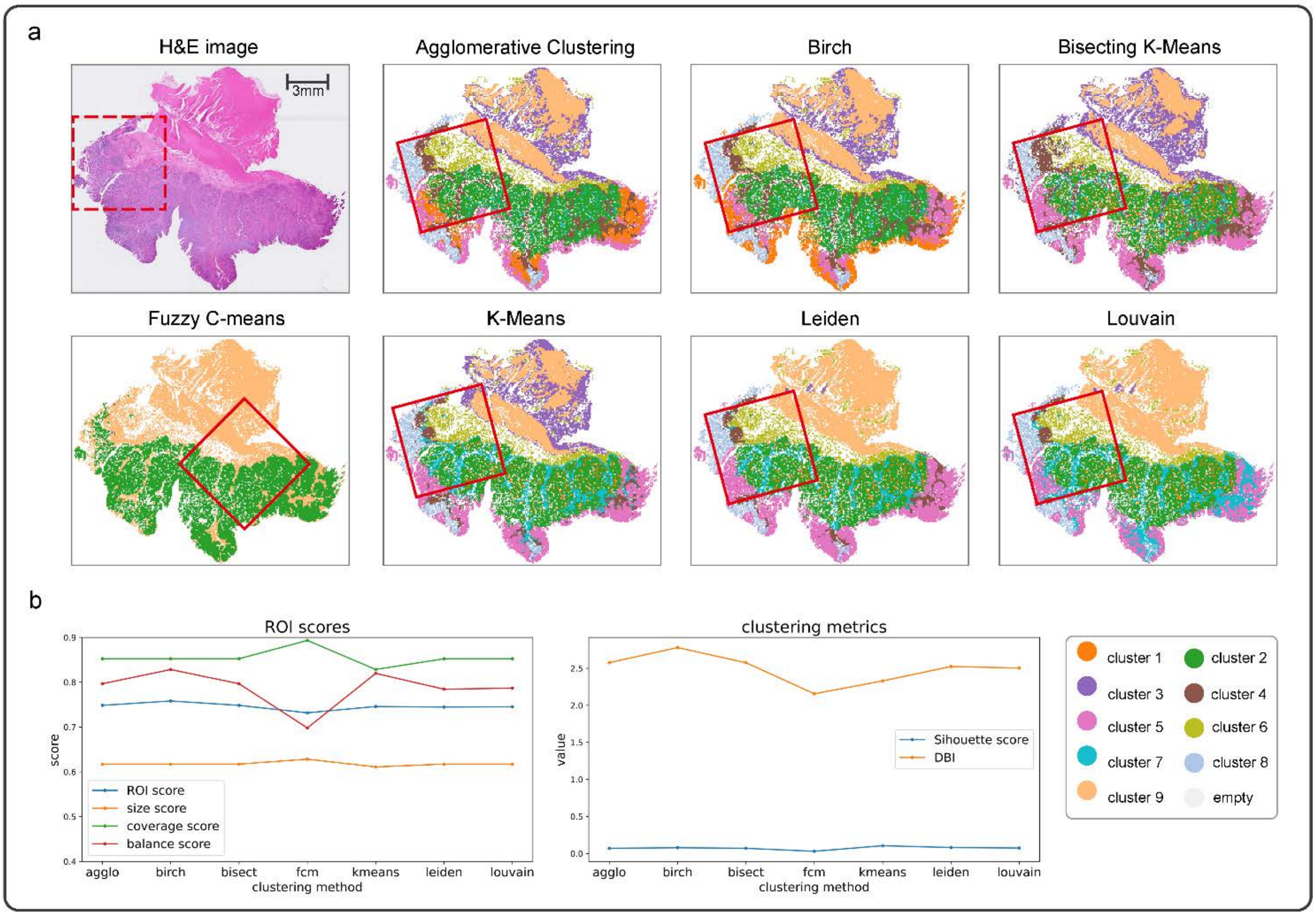

**Supplementary Fig. 23 | Composition of ROI score.** ROI score is the geometric average of balance score, coverage score and size score. All three sub-scores range from 0 to 1. The more similar the distribution of histology clusters inside the ROI is to uniform distribution, the larger the balance score. The less empty region a ROI includes, the larger its coverage score. The bigger the size of ROIs, the larger the size score.

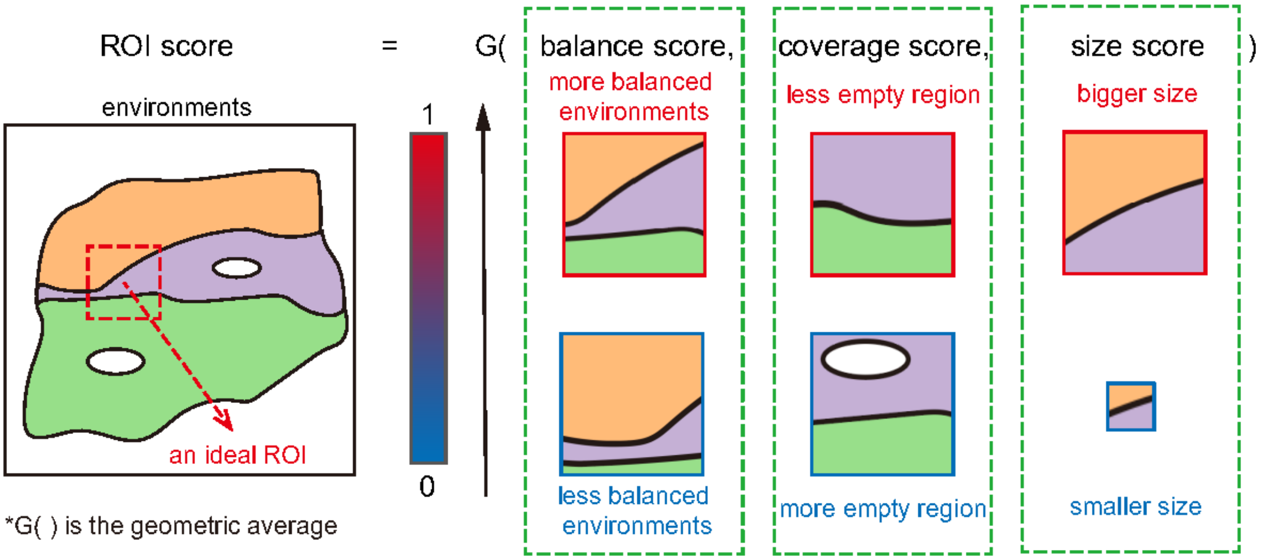
